# Human Bone Marrow Plasma Cell Atlas: Maturation and Survival Pathways Unraveled by Single Cell Analyses

**DOI:** 10.1101/2023.01.18.524601

**Authors:** Meixue Duan, Doan C. Nguyen, Chester J. Joyner, Celia L. Saney, Christopher M. Tipton, Joel Andrews, Sagar Lonial, Caroline Kim, Ian Hentenaar, Astrid Kosters, Eliver Ghosn, Annette Jackson, Stuart Knechtle, Stalinraja Maruthamuthu, Sindhu Chandran, Tom Martin, Raja Rajalingam, Flavio Vincenti, Cynthia Breeden, Ignacio Sanz, Greg Gibson, F. Eun-Hyung Lee

**Author notes:** Senior Co-authors (Greg Gibson, F. Eun-Hyung Lee). Authors contributed equally as the second authors (Doan C. Nguyen, Chester J. Joyner). Corresponding author: F. Eun-Hyung Lee, MD, Division of Pulmonary, Allergy, Critical Care & Sleep Medicine Emory University 615 Michael Street, Suite 205 Atlanta, GA 30322.

## Abstract

Human bone marrow (BM) plasma cells are heterogeneous, ranging from newly arrived antibody-secreting cells (ASC) to long-lived plasma cells (LLPC). We provide single cell transcriptional resolution of 17,347 BM ASC from 5 healthy adults. Fifteen clusters were identified ranging from newly minted ASC (cluster 1) expressing MKI67 and high MHC Class II that progressed to late clusters 5-8 through intermediate clusters 2-4. Additional clusters included early and late IgM-predominant ASC of likely extra-follicular origin; IFN-responsive; and high mitochondrial activity ASC. Late ASCs were distinguished by differences in G2M checkpoints, MTOR signaling, distinct metabolic pathways, CD38 expression, and utilization of TNF-receptor superfamily members. They mature through two distinct paths differentiated by the degree of TNF signaling through NFKB. This study provides the first single cell resolution atlas and molecular roadmap of LLPC maturation, thereby providing insight into differentiation trajectories and molecular regulation of these essential processes in the human BM microniche. This information enables investigation of the origin of protective and pathogenic antibodies in multiple diseases and development of new strategies targeted to the enhancement or depletion of the corresponding ASC.

**One Sentence Summary:** The single cell transcriptomic atlas of human bone marrow plasma cell heterogeneity shows maturation of class-switched early and late subsets, specific IgM and Interferon-driven clusters, and unique heterogeneity of the late subsets which encompass the long-lived plasma cells.

## INTRODUCTION

The existence of long-lived plasma cells (LLPC) that provide a lifetime of humoral protection after vaccination and infection is well-established in mice and humans. Activated lymph node B cells differentiate into early antibody secreting cells (ASC) which home to bone marrow (BM) niches where a fraction may survive as LLPC, which we previously identified within the CD19^-^ CD38^hi^CD138^+^ cell population (*1–3*) (*4–6*). Yet, the complexity of this population has never been investigated and thus, the precise phenotype of LLPC, their differentiation pathways and survival programs need to be understood. In addition, whether mere migration to protective niches is sufficient for the establishment of a LLPC compartment or instead, additional maturation in the BM is required, remains to be elucidated. We recently reported that early maturation of nascent ASC takes place in the BM through morphologic, transcriptomic, and epigenomic changes that presumably enable their ultimate differentiation into LLPC (*6*). Accordingly, we postulated that peripheral ASC arriving in the marrow must comprise needs to mature locally to generate *bona fide* LLPC.

We have recently developed a novel *in vitro* human BM mimetic system, containing soluble factors from mesenchymal stromal cells (MSC), APRIL, and hypoxia, which sustains ASC survival for up to 56 days in culture, thereby overcoming previous experimental limitations in the field imposed by the rarity and *ex vivo* frailty of ASC. This approach enabled us to generate a molecular roadmap charting the progression of nascent ASC into mature ASC demarcated by upregulation of CD138 expression and downregulation of CD19 as early as day 14 (*7*). This early work showed that engagement of sequential transcriptomic and epigenetic programs promoting resistance to apoptosis were essential for survival (*6*). Although revealing of candidate LLPC BM maturation programs, these bulk culture analyses could not directly interrogate the BM LLPC compartment.

To address this unmet need, we have now performed extensive single cell analysis of BM ASC. Our studies identify a large degree of cell heterogeneity and maturation trajectories. Starting from nascent KI67+ ASC with the highest MHC Class II expression, we could assign early and late ASC distinguished by differences in G2M check points, E2F targets, MTOR signaling, and metabolic pathways for fatty acid and oxidative phosphorylation. Early and late BM ASC also displayed differential expression of members of the TNF receptor superfamily and CD38 expression. Finally, terminal differentiation of late IgG ASC followed two distinct paths with differential utilization of the TNFα signaling pathway via NFkappaB. In all, this study provides the first single cell resolution cell atlas and molecular roadmap trajectories of LLPC maturation in the human BM.

## RESULTS

### Single-cell transcriptomic profiling of human plasma cells in the BM

To characterize the heterogeneity of human bone marrow plasma cells (BMPC), the three major populations previously identified were sorted from healthy adults without recent immunization or infection (*3*): pop A (CD19^+^CD38^hi^CD138^-^), pop B (CD19^+^CD38^hi^CD138^+^) and pop D (CD19^-^ CD38^hi^CD138^+^) (Fig. 1A). We sought to test the hypothesis that while our early studies indicate that Pop D constitutes the main reservoir of LLPC, this population is nonetheless heterogeneous and contains different subsets with distinct molecular programs reflecting their generation, regulation, and survival. We tested this hypothesis by studying BM ASC at the single cell level using the 10x Genomics Chromium platform in conjunction with 5’ directed cDNA library construction to characterize both the single-cell transcriptomic profiles (scRNA-seq) and matching single-cell V(D)J repertoire sequencing (scVDJ-seq). After exclusion of non-ASC contaminating cells (including B cells), low-quality cells, dying cells and doublets, the remaining cells were identified as *bona fide* ASC with characteristic expression of *XBP1*, *IRF4*, *PRDM1*, *CD27*, *CD138*, and *CD19* (Fig. 1B, C, E and Fig. S1C, S2A). In total, we retained 17,347 ASC for processing through our scRNA-seq analytical pipeline.

**Figure 1.**
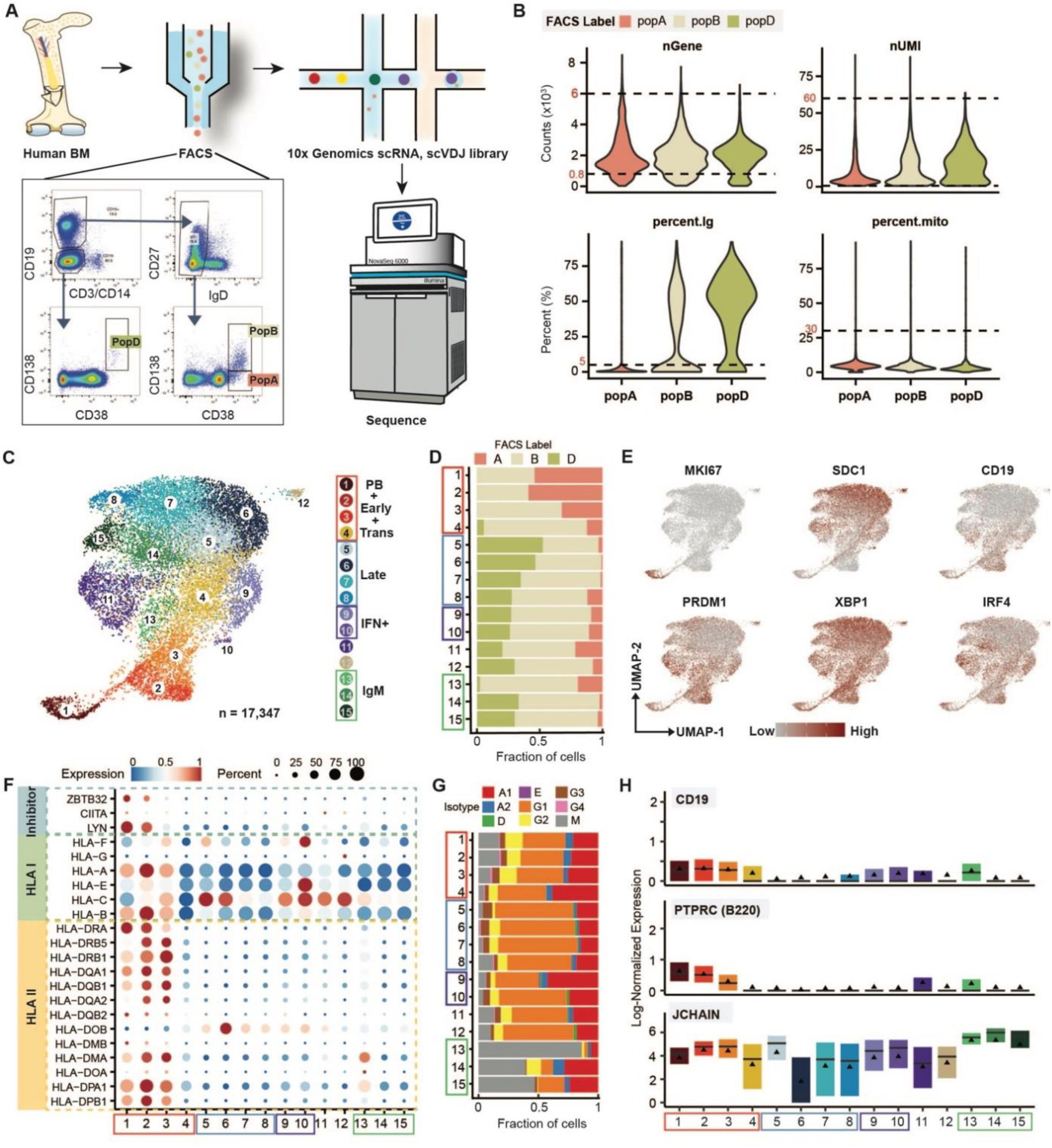
Single cell Transcriptomic Profiling of bone marrow plasma cells. A) Schematic of single-cell RNA profiling for human BMPC. B) Criterion for removing bad quality cells. Dashed lines show the cutoffs which are labeled in red. C) scRNA-seq cell clusters of combined data from 5 healthy bone marrow patients and visualized in UMAP colored by cell types. Red and blue boxes highlight the early and late stage of BMPC maturation, respectively. The purple box highlights the path towards IFN response PC subgroups and the green box highlights IgM dominant cell populations. D) The fraction of cells from FACS-sorted cell population in each cell subgroup identified in C). E) Key PC-associated master gene expression. The redder the dot indicates the higher log-normalized gene expression. F) Dot plot for expression of human MHC class I, II and inhibitors of class II genes. Here and in later figures, colors represent Min-Max normalized mean expression of marker genes in each cell group, and sizes indicate the proportion of cells expressing marker genes. G) The fraction of isotypes identified by scVDJ-seq data in each cell subgroup identified in C). H) Boxplots showing the expression levels of the indicated genes. The solid triangle represents the average value.

### Identification and ordering of 15 clusters of BM ASC by single cell profiling

Clustering was performed with removal of known Ig genes and without knowledge of subsets A, B, and D a priori. However, our initial clustering required elimination of two subsets falsely identified solely on the basis of very high expression of two misannotated Ig VH genes (Materials and Methods). Ultimately, we identified 15 robust clusters representing different cell states or types based on gene expression profiles identified with the canonical correlation analysis workflow in Seurat (Fig. 1C, E and Figs. S2A, B, C, see Materials and Methods). Accordingly, Rand Index (RI) (*8*), which calculates the concordance of pairwise relationships between all pairs of cells, typically results in 86.5% similarity on average in cluster designation for each individual cell (Fig. S2D, see Materials and Methods). Notably, the 15 subgroups were detected in all 5 individuals, with no identifiable subject-specific variability or Ig driven clusters (Figs. S2C,E,G). After establishing the 15 clusters, we re-incorporated the Ig genes and found no particular cluster with a specific VH gene family driving a BM ASC cluster. Finally, the 15 distinct clusters (Fig. 1C) could be grouped as early (clusters c1-3); transitional or intermediate (c4); and late (c5-8) ASC based on HLA class I and II expression (Fig. 1F). In addition, we identified 2 IFN-response dominated clusters (c9-10); and 3 IgM-predominant clusters (c13-15, Fig. 1G), that contained the majority of BM IgM ASC. Interestingly, c13 shared most features of early clusters while c14-15 displayed late features, thereby suggesting the IgM counterpart of early and late IgG clusters. Finally, we identified a mitochondrial-high cluster (c11) with characteristics of early ASC and a minor cluster (c12), likely representing dying ASC. The differentially expressed marker genes used to adjudicate each of the clusters are shown in Fig. S2F and Table 1. Their distinctive transcriptional profiles, transcription factors networks, characteristics of antibody repertoires and inter-relationships within differentiation trajectories are discussed in detail in the following sections.

**Table 1.** Information of cell clusters defined in Fig.1C

| Cluster id | Cell label | Markers | Cluster id | Cell label | Markers |
| --- | --- | --- | --- | --- | --- |
| 1 | PB | MKI67, UBE2C, BIRC5, SPN, TUBB | 9 | pc9 | ATF5, PSAT1, CEBPB, ERN1 |
| 2 | Early1 | CD52, PEBP1, MHC Class II (except HLA-DOB) | 10 | IFN+ | MX1, XAF1, IRF7, STAT1 |
| 3 | Early2 | CD79A, CD74 | 11 | Mito-high | MALAT1, IRF4, PRDM1, ZBTB20, MTRNR2L12, FCRL5 |
| 4 | Trans | IGLV3-25 | 12 | pc12 | JUND, CXCR4, SIK1B |
| 5 | Late1 | SMOC1, MOXD1 | 13 | IgM1 | CCDC88A, FOXP1, IGHM, KLHL14, MS4A1 |
| 6 | Late2 | CD9, CST3, CD63, TIMP1 | 14 | IgM2 | JCHAIN, RGS2, TNFRSF4 |
| 7 | Late3 | RHOB, CDKN1A, RGCC | 15 | IgM3 | CCL3, CCL4 |
| 8 | Late4 | JUNB, FOS, EGR1, NR4A1 |  |  |  |

### Separation of early and late clusters by HLA class II gene expression

Cluster 1 distinctly expressed MKI67, CD43 (*SPN*) and CD27, and had low expression of CD20 (*MS4A1*, Figs. S2F, G). This pattern is consistent with the definition of proliferative plasmablasts (PB) or early minted ASC, likely representing new BM arrivals from active immune responses (CD20^low^CD27^high^CD43^high^) (*9*). As they mature into resting LLPC, proliferative ASC typically shed B cell markers such as surface Ig and MHC class II genes (encoded by human leukocyte antigens gene complex (HLA))(*3, 4*). Clusters 2 and 3 lacked Ki67 expression but retained the highest levels of HLA class II genes, which subsequently decreased in the transitional cluster 4 (Fig.1F). Thus, the initial ASC seeding the BM could be further divided into early stages including proliferative and non-proliferative cells (c1 and c2-3, respectively); and transitional stages (cluster 4). Interestingly, c1-3 were devoid of Pop D, which could be first detected at low frequencies within cluster 4 (Fig. 1D).

As expected, *PRDM1,* encoding BLIMP1 which drives PC commitment and extinguishes class II transactivator *CIITA* and MHC class II gene expression, was expressed from the early ASC stages (*10*) (Fig. 1F). In turn, *CIITA* was universally extinguished on nearly all BM ASC subsets except for a small fraction of early c1. Since *CIITA* is a master regulator of MHC class I and II genes, its expression pattern is consistent with the predictable downregulation of HLA in mature BM ASC subsets. On the other hand, HLA-DOB, which suppresses peptide loading of class II molecules, was upregulated in the late ASC clusters which is consistent with reports that HLA-DOB is less affected by *CIITA* than other family members (*11*). HLA class I expression followed similar trends except for HLA-C. In all, HLA class II expression provides clear separation of early and late BM ASC subsets.

### Transcriptional characteristics of human BM ASC clusters

We next identified cell type-specific markers across the 15 BM ASC subgroups (summarized in Fig.S2F, Table S1, Materials and Methods). In addition to the defining features, Ki-67 and HLA class II, other markers of early ASC included *LYN* and *ZBTB32*. LYN is a tyrosine protein kinase downstream of BCR signaling that can diminish proliferation while driving terminal PC differentiation (*12*) (Fig.1F). Interestingly, LYN deficiency leads to a 20-fold increase of BM PC and induces autoimmune disease in mouse models (*13*). Thus, progressive extinction of LYN expression in late clusters may contribute to enhanced survival of the more mature PC(*14*). Also informative for early maturation was the expression of B cell markers *CD19* and *PTPRC* (B220). Similar to *CD19*, *PTPRC* was strongly expressed in early IgG-dominant clusters 1-3 as well as in cluster 11 and IgM-dominated cluster 13 (Figs. 1G, H). The progressive loss of *CD19* and *PTPRC* during B cell differentiation into PC and the ability of CD19+ B220+ cells to secrete low-affinity IgM antibodies (*15, 16*), also supports our inclusions of clusters into early and late populations.

The transitional nature of cluster 4 was documented by the progressive loss of early features including MHC Class II and initial expression of late markers. In turn, c5 shared with c4 higher expression of lambda light chain V3-25 while displaying higher levels of transcripts characteristic of late subsets including *MDK*, *ITM2B*, *LMNA*, *AREG*, and *TIMP1* (Table 1 and Table S1).

Late clusters 6-7 and to a lesser extent c8, expressed higher levels of *CD9* and *CST3*. The CD9 tetraspanin modifies multiple cellular events of relevance for BM ASC including adhesion, migration, proliferation, and survival. *CD9* expression has been considered to mark GC-derived human plasma cells (*17*); however, *CD9* is also considered a marker of mouse PC derived from marginal zone and B1 B cell in primary T-dependent responses (*18*)

Finally, genes involved in regulation of cell cycle, proliferation, apoptosis, and survival were preferentially expressed in clusters 7 (*CDKN1A*/p21 and *RGCC*), and 8 (*JUNB*, *FOS*, *EGR1*, *NR4A1*) (Table 1). P21 is a potent cyclin-dependent kinase inhibitor whose expression regulates cell cycle progression at G1 and is tightly controlled by the tumor suppressor p53 which mediates cell cycle arrest, can promote apoptosis in a context-dependent fashion, and induces *RGCC* which also regulates cell cycle progression (*19*). Early Growth Response (*EGR1*) is a nuclear transcriptional regulator of multiple tumor suppressors including p53. Notably, *EGR1* is rapidly induced by growth factors, apoptotic signals and hypoxia, a feature of the BM microenvironment that determines ASC survival (*20, 21*). Of note, it has been shown to play a non-redundant role in PC differentiation (*22*). While the induction of the orphan nuclear receptor *NR4A1* (Nur77), is best recognized as a consequence of antigen receptor engagement in B and T cells (*23*), *NR4A1* is also induced by other stimuli including ER stress which is present at high levels in PC secondary to high immunoglobulin synthesis (*24*). Interestingly, *NR4A1* can modify the pro/anti-apoptotic balance of the Bcl2 family and has enhanced binding to anti-apoptotic Bcl-B which is prominently expressed in PC (*25, 26*).

Notably, clusters 7 and 8 separated from other late ASC populations (c5 and c6), by the highest levels of the TNFα-signaling NFκB pathway. These features, the transcription factor regulatory networks and their potential participation in separate maturation trajectories are presented in detail in subsequent sections. Suffice it to say at this time that IFN responsive genes (IFN+) were the major markers in clusters 9 and 10. Cluster 9 highly expressed *ATF5, CEBPB* and *ERN1* whereas cluster 10 expressed the classical IFN response signature (*STAT1, IRF7, ISG15, IFITM1, IFI6, MX1* and *OAS1*). Given the prominent role that Type I and Type II IFN responses may play in lupus, it will be important to determine whether these clusters are preferentially expanded in this autoimmune disease (*27*).

We identified all five isotypes (IgM, IgG, IgA and a small fraction of IgD and IgE), as well as the four IgG subclasses (IgG1-IgG4), in BM ASC (Fig. 1G). The majority of ASC clusters were dominated by IgG, predominantly IgG1. Compared with early stages, late stages (clusters 5-8, and 10-12) had expanded proportions of the IgG isotype (p value _one-sided proportional test_ < 2.2e^-16^). Clusters 13-15 contained the largest repository of IgM cells (40-80%), which represented a large majority of cluster 13. These clusters are candidates for the human counterpart to mouse IgM LLPC that accumulate in the spleen in a GC-independent fashion and contribute to protective IgM responses (*28–30*).

The separation of clusters 13-15 was driven by their over-expression of the IgM isotypes, representing 40-85% of cells in these populations (Fig. 1G). Interestingly, these IgM-predominant clusters displayed an overall distinct transcriptome and could in turn be split into early (c13) and late clusters (c14-15), according to the features described above for IgG-dominant clusters. This determination is also consistent with the scarcity of Pop D cells in cluster 13, whereas clusters 14 and 15 contained a frequency of Pop D similar to late IgG clusters 5-8 and a virtual absence of Pop A (Fig. 1D). Combined, these observations define clusters 1-3 and 5-8 collectively as early and late phase/stage of PC maturation respectively, and clusters 9-10 and 13-15 collectively as IFN+ driven and IgM dominant clusters, respectively (Fig. 1C, Fig. S2C). Of interest, J-chain transcription, typically ascribed to dimeric/polymeric IgA and IgM-producing cells, was significantly active across all BM ASC clusters, albeit at higher level in IgM-dominant c13-15. These findings are consistent with previous data indicating J-chain transcription in different stages of B cell differentiation (*31*).

### Identification of mature IgM ASC populations in the human BM

IgM ASC (clusters 13-15) are further defined by higher expression of *CCR10*, *JCHAIN*, *FHL1*, *PHACTR1* and *RAMP2* relative to IgG dominant cell populations (Fig. S3A, Fig. 1H). Since C-C motif receptor 10 (CCR10) and JCHAIN are widely expressed in mucosal ASC (*31, 32*), the IgM dominant cell populations may have mucosal origins. Additional DEG distinguishing IgM from IgG dominant clusters included 77 *CCR10* co-expressed genes summarized in Fig. S3B. These *CCR10* related genes included *EBI2* (*GPR183*), *JCHAIN, FOXP1*, and several genes involved in regulation of lymphocyte activation such as *FCRL3, TNFSF9, CLECL1* and *FGL2*. *FOXP1* is known to impair the formation of germinal centers (GC) and to repress human plasma cell differentiation but is highly expressed in mature primary human B cells (eg. Naïve B, memory B cell) as well as in mouse follicular B and B-1 cells (*33, 34*). This profile is consistent with a GC-independent extrafollicular origin of the late IgM ASC (*34, 35*).

### Relationship of BM ASC single cell clusters to previously described ASC populations

We sought to delineate the transcriptional levels of surface markers previously used to define different BM ASC populations including the Pop D containing the LLPC of greatest longevity(*3*). Consistent with previous models of BM ASC maturation, *CD19* expression was notable in early clusters 1-4 and extinguished in late clusters while SDC1 (*CD138*) expression was highest in the late clusters (Fig. 1E). *XBP1*, an essential transcription factor (TF) associated with the ASC unfolded protein response (UPR), was increased in all clusters. Contrary to mouse studies, *PRDM1* expression was higher in early subsets and did not continuously increase in all late subsets (Fig. 1E, Fig. S4A). Thus, *PRDM1* upregulation was limited to a small fraction of late clusters 7, 8 and 15, indicating that the activity of this essential plasma cell transcription factor may be required for early ASC differentiation yet less important for terminal differentiation or maturation and long-term survival (*36, 37*).

The majority of Pop D was found in the late stages of ASC maturation and interestingly, it was distributed across clusters 5-8, which also included a minority of pop A cells but transcriptionally resembled pop B (Fig.1D, Fig. S4D). A smaller fraction of Pop D was spread across late clusters 5-12 and 14-15. In contrast, pop B contributed nearly half of the cells in each of the early and late clusters, again suggesting that it corresponds to an intermediary BM population. However, it is interesting that pop B in the late cluster 5-8 had very few cells expressing CD19 suggesting that surface protein was lagging RNA expression. Combined, these findings demonstrate substantial heterogeneity of mature BM ASC and suggest that ASC of LLPC potential may be present in several BM compartments.

By contrast with the late clusters, clusters 1-4 were enriched for pop A ASC and devoid of pop D (Fig. 1E). Both these early and a “mito-high” subgroup (c11), had higher *CD19* and *PRDM1* expression, higher number of total detected genes and lower percentage of Ig transcripts relative to late mature ASC (Fig. 1E, Figs. S4A, B, C). Combined with their higher expression of *IRF4*, *ZBTB20*, and *FCRL5* (Table 1), this cluster appears to belong with the early ASC populations (*38*).

### Identification of 4 maturation paths for IgG BM ASC through trajectory analysis

IgG ASC comprise the major isotype in the late clusters, thus, we used Slingshot to construct maturation trajectories to focus on IgG BM ASC and visualize the dynamic alteration of gene sets (Fig. 2A, Materials and Methods). This approach predicted five maturation trajectories. Path1 and 2 projects to late clusters 6 and clusters 7 & 8 respectively. Since path3a and path3b direct to the same terminal cluster 9, we merged them into a single path3 (Fig. 2A). Finally, path4 leads toward cluster 11. We projected the pseudotime for each path onto the UMAP clusters (Fig. 2B). There was distinct separation between clusters 1 and 2; but the remaining non-proliferative clusters in each path tend to be shared between adjacent stages/clusters. Once the short-lived early ASC pass through the proliferative stage, the maturation of BM ASC is more of a continuum of functional processes rather than an ordered sequence of discrete cell states.

**Figure 2.**
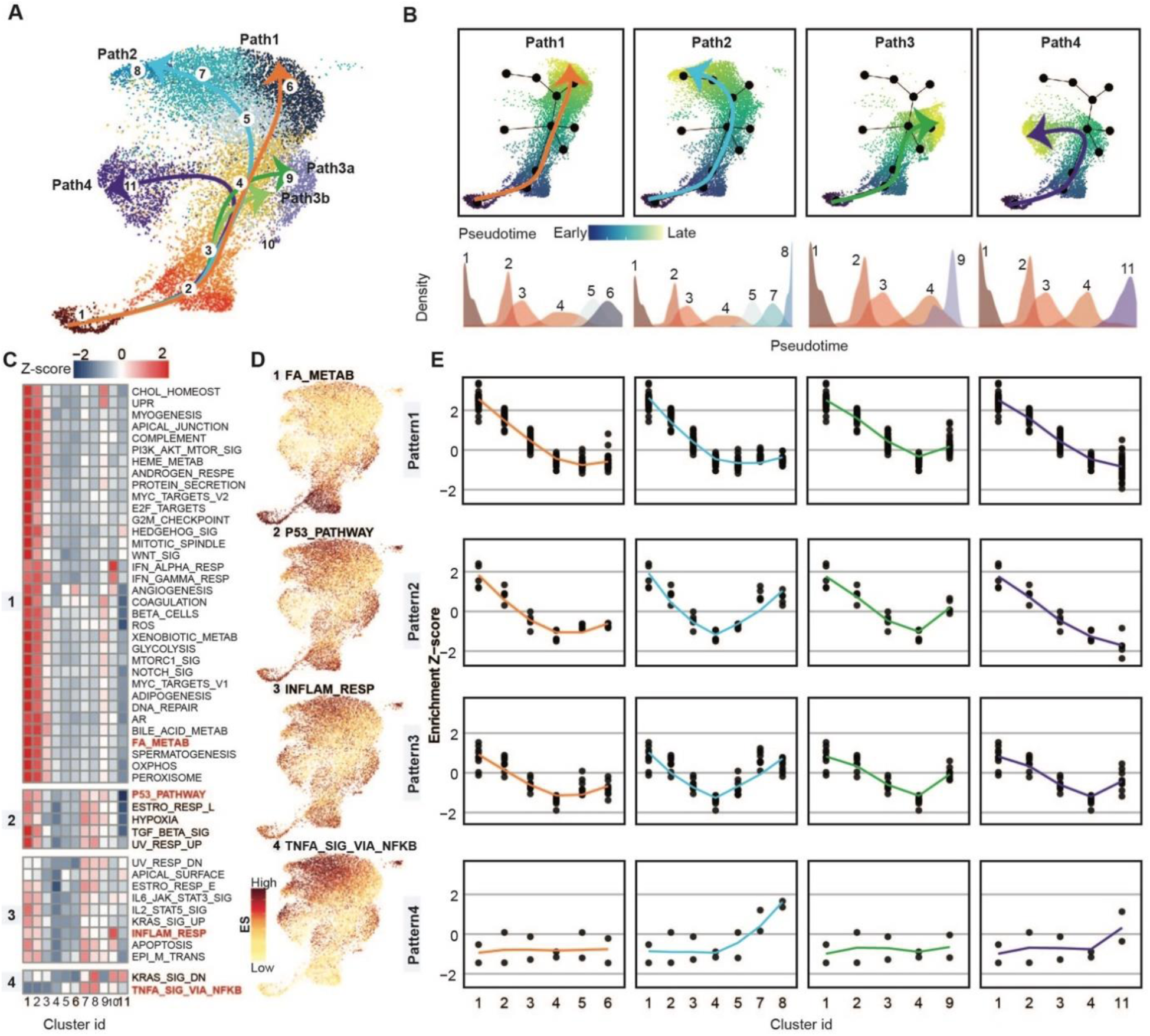
Trajectory and Hallmark pathway enrichment analysis to distinguish predicted BMPC maturation paths. A) UMAP plot shows predicted paths of IgG1 dominant BMPC maturation. Arrows indicate the maturation direction. B) UMAP plots show cell populations located in each maturation path and colored by predicted pseudotime (top), each solid black dot represents a cell population used to predict path in (A). The darker the blue and the lighter the green indicate the earlier and later stages of maturation, respectively; (bottom) The density plots show the distribution of scRNA-seq identified cell subgroups on pseudotime space of each path. C) Heatmap shows the row-scaled enrichment scores (ES) for Hallmark pathway enrichment analysis in the IgG1 dominant cell populations, from left to right is cluster 1 to 11. The space in the heatmap indicates the boundaries of different patterns detected by two-way hierarchical clustering in all the cell populations (Fig.S8). The pathways labeled in red are selected examples from each pattern for visualization in (D). D) Projected the enrichment score of indicated pathways from each pattern onto UMAP. The darker the red, the higher the enrichment. E) Dotplot next to the example umap visualization in (D) showed the scaled enrichment scores from each pattern in cell populations from each path. X-axis is ordered cell populations corresponding to the cell order in each path in (B). “Loess” method fitted lines of ES alteration trends were colored by predicted paths in (B).

Next, Hallmark pathway enrichment scores (*39*) were used to gain insight into the gene activity distinguishing the four branches of IgG BM ASC maturation (see Materials and Methods). In total, we discovered 4 patterns describing the dynamic alteration of gene sets during BM ASC maturation (Fig. 2C, Fig. S5A). From each pattern, a pathway example for visualization is shown in Fig. 2D. Projecting scaled enrichment scores from each pattern in Fig.2C onto pseudo-space paths inferred in Fig. 2B, patterns1, 2, and 3 exhibit a linear-like decrease in the pre-late stages but slightly different enrichment patterns in the late-phase (Fig. 2E). More specifically, pattern1, which includes pathways of the unfolded protein response (UPR), reactive oxygen species (ROS), oxidative phosphorylation (Oxphos), and fatty acid metabolism (FA_METAB), decreases but then plateaus in late-phase of maturation for paths1, 2, and 3. Only path4 decreases continuously into the late phase (Fig. 2E). Pattern2 includes the hallmark pathways: hypoxia, UV response up and p53 pathway showing the same maturation trends but an increase in late states of path2 corresponding to clusters 7 and 8. Pattern3 contains IL6-JAK-STAT3 signaling, inflammatory response and apoptosis revealing the similar maturation trends in all 4 paths. Pattern4 notably including TNFα signaling via NFκB and KRAS signaling down is most intriguing since the enrichment diverges between the late ASC in path1 and 2. In addition to the IgG trajectories, we found higher enrichment score in IgM-pre-dominant lineage (c13-15) for TNFa signaling via NFκB (Fig. S5B), but it also showed an increasing trend during the maturation of IgM-dominant lineage (Fig. S5C).

### Transcription factors of ASC maturation and survival

To understand distinct enrichment patterns of gene sets regulated by specific transcription factors (TFs), we imputed the potential functions of TFs in regulating BM ASC maturation (see Materials and Methods). We observed 205 differentially expressed TFs that are expressed in at least 10% of assigned cell clusters (see Materials and Methods and Table S2). For visualization, the most abundant 144 TFs were assigned to the cell cluster or defined stage of maturation (Fig. 3A, Fig. S2C) with the highest gene expression. These “cluster-associated” or “stage-associated” TFs, were observed in 9 out of 15 cell groups. Notably absent were any TFs defining the transitional and late-stage clusters 4-7 or 14 and 15.

**Figure 3.**
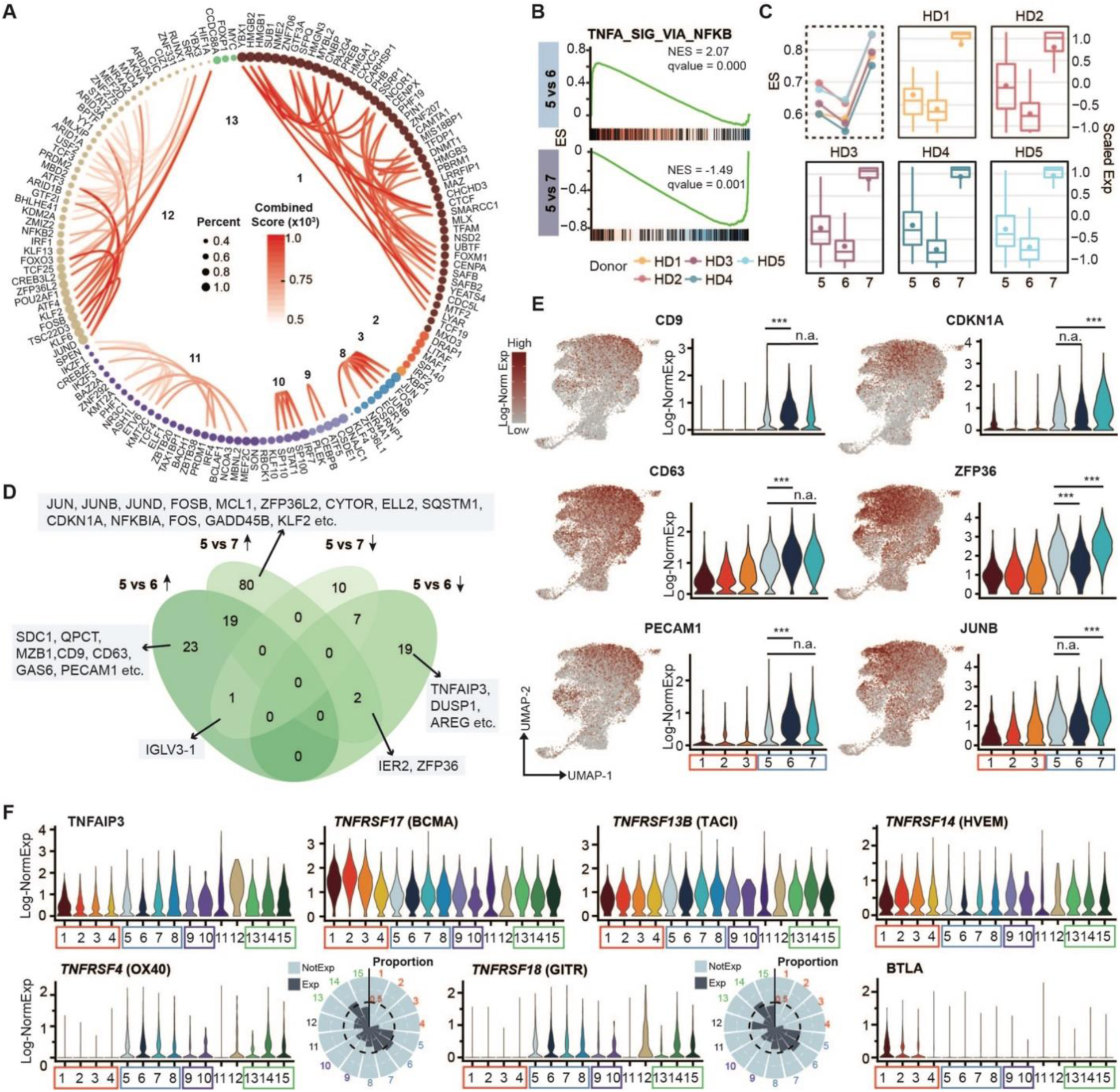
Genes and Hallmark pathways distinguish late phase maturation fate. A) Cell cluster associated TFs, each node represents a TF, colored by associated cell cluster and scaled by expressed percentage of cells in the cluster. The lines between nodes are inferred protein-protein interactions from the STRING database. The redder the line, the more confident the inferred interactions. Numbers in the circles show the assigned cell cluster id. B) GSEA of the most indicated pathways TNFα singaling via NFκB that are differentially enriched between 5 vs 6 and 5 vs 7. C) TNFα singaling via NFκB hallmark pathway ES in 5 subjects (in dashed box and y-axis is on the left) and the distribution of scaled corresponding pathway enriched maturation-associated DEG expression in cluster 5, 6 and 7 in each individual (in solid boxes and y-axis is on the right). D) Comparison of top DEGs between cluster 5 vs 6 and cluster 5 vs 7 (see Methods), based on the sign of avglogFC, the DEGs were divided into up-/down-regulated in cluster 6 or 7 groups. Venn diagram shows the results of the DEG comparison. E) Gene expression examples from the results of the comparisons in (D). F) Gene expression of indicative genes from TNF family. X-axis is cell cluster id and y-axis is the log-Normalized gene expression. Circular barplot showed the proportion of cells in each cell subgroup showed expression of corresponding genes. Black dashed line indicates the proportion of 0.5 and numbers indicate the cell cluster ids. NotExp: Not expressed; Exp: expressed; *** indicates Bonferroni adjusted p value < 0.001; n.a.: not available.

Adding protein-protein interaction information from the STRING database (*40*), we observed that some TFs are cluster-specific and/or stage-specific (early or late) indicating that TFs may regulate associated functions in cluster-specific or stage-specific ways (Table S2). Thus, 48 out of 144 (33%) of the TFs were cluster 1-associated, and typically known to be involved in regulating basic functions such as RNA/DNA binding, transcription, or metabolic processes, RNA stabilization (such as YBX1, SUB1), high-mobility group protein B (HMGB) damage-associated molecular patterns (DAMPs)(*41*), (such as HMGB1, HMGB2 and HMGB3) and cell cycle progression (such as MYBL2(*42*)) among others (eg. GTF3A, HMGN3, SPI1). Similarly, TF important for immune activation, signaling, differentiation and survival (EGR1, JUN, JUNB, FOS, NR4A1 and ZFP36L1), were cluster 8-specific rather than stage-specific.

Two other large groups of 25 (∼17%) and 43 (∼30%) TFs were identified in cluster 11 and cluster 12, respectively. While high mitochondrial gene expression suggests Cluster 11 cells may be experiencing high stress or be destined for death, they were nevertheless characterized by high expression of *IRF4*, *PRDM1*, *ZBTB20*, *IKZF3* (AIOLOS protein) and *IKZF1* (IKAROS). This profile is also consistent with a GC-derived BM ASC subset with enhanced survival that is different from the other late IgG subsets (*29, 43*). Previous studies have shown that the absence of AIOLOS affects LLPC survival as well as the production of high-affinity antibodies but does not perturb ASC homing (*44, 45*), indicating an important role in support of the survival of high-affinity antigen-specific LLPC. IRF4 is required for both activation of B cell and GC formation, and it initiates PC differentiation in a dose-dependent manner (*46–49*). High levels of IRF4 result in repression of BCL6 and activation of the PC transcription factor BLIMP1 as well as the BCL-6-related transcription factor ZBTB20, which may be related to PC differentiation, survival and longevity (*36, 38, 47*). Consequently, survival fate decisions may be occurring in cluster 11. Cluster 12 was characterized by accumulation of ATF4, the first stable product that can trigger the activation of the PERK pathway, which promotes ASC apoptosis when ER stress is unabated (*50, 51*). Thus, this small cluster may indeed be a death-prone ASC.

In contrast to cluster-specific TF, *XBP1* showed increased expression in the early phase of ASC maturation, with the highest gene expression in cluster 3. *XBP1* is involved in protein glycosylation and lipid synthesis and is a component of the IRE1 branch of the UPR which plays a role in elevating ASC endoplasmic reticulum (ER) capacity. Interestingly, the highest *XBP1* expression occurred in the early stages suggesting its potential importance in accommodating ER expansion and cell survival in the initial phase of BM ASC maturation.

### Bifurcation of LLPC differentiation

To characterize the gene sets involved in the bifurcation of late IgG ASC maturation in path1 and 2, we examined the differentially enriched hallmark pathways between clusters 5 vs 6 and clusters 5 vs 7 using GSEA prerank analysis (see Materials and Methods). GSEA prerank analysis identified 6 and 19 gene sets significantly activated or repressed with FDR < 0.05 during BM ASC maturation from cluster 5 to 6 and 5 to 7, respectively (Table S3). Of these, 5 gene sets exhibited the opposite direction of enrichment in the two paths (one activated and the other repressed) while 16 showed the same directional trend. Although IFN-γ response and allograft rejection were downregulated in cluster 6, inflammation and MYC targets v1 were upregulated in cluster 7, only TNFα signaling via NFκB was significant for downregulation in c6 and upregulation in c7 (FDR ≤ 0.001) (Fig. 3B). To further confirm that the enrichment difference in TNFα signaling via NFκB was not driven by any one subject, we evaluated the pathway enrichment score (ES) in associated clusters for each individual. In all 5 individuals, ES was downregulated in cluster 6 but upregulated in cluster 7 (Fig. 3C). Thus, this conclusion is not driven by any one individual or any outlier genes but is rather a general trend (see Fig. 3C, Materials and Methods).

Furthermore, by comparing gene expression between cluster 5 vs 6 and cluster 5 vs 7, the Venn diagram in Fig. 3D picked top DEGs and divided associated top DEGs into cluster 6 or 7 up-/down-regulated gene lists specific and common to each cluster (Table S4, see Materials and Methods). The expression of corresponding genes is visualized in Fig. 3E.

Subsequently, we compared the TNFα signaling via NFκB directly between cluster 6 and 7 and discovered 78 genes contributing to the significant enrichment (Fig. S6A,B, Table S5). Out of those 78 leading edge genes, thirty-four (∼44%) are among the top DEGs including *TNFAIP3, FOS, NFKBIA, CDKN1A*, and *ZFP36* (Fig. S6C, Table S4, Table S5, Materials and Methods). Interestingly, path2 had mildly increased *PRDM1* expression suggesting upregulation in this late cluster only (Fig. S2D).

### Differentially expressed TNFRSF family members between early and late ASC

TNF and TNF superfamily cytokine signaling play important roles in B cells and plasma cell survival and function or cell death. The best known factors BAFF or APRIL (*TNFSF13*) and their recognized receptors BAFF-receptor (BAFFR:*TNFRSF13C*), Transmembrane Activator and CAML-interactor (TACI:*TNFRSF13B*) and B-cell maturation antigen (BCMA:*TNFRSF13A/17*) are essential for ASC survival (*52, 53*). APRIL binds strongly to BCMA and moderately to TACI whereas BAFF binds weakly to BCMA and strongly to TACI (*54*). Although BAFF has been suggested to be important in mouse BM ASC (*55*), APRIL is critical for ASC survival and maturation in both mouse and human (*7, 56*). Finally, APRIL and not BAFF can bind to heparin sulfate proteoglycans (HSPGs) such as CD138 which likely concentrates APRIL at the cell surface thereby increasing its effectiveness (*57*).

We found little difference in expression of the members of the TNFRSF family between the two late IgG paths1 & 2; however, there were major differences between the early and late clusters (Fig. 3F, Figs. S7A, B). BCMA showed high expression in early ASC and was significantly downregulated in late ASC clusters while TACI expression was significantly increased in late clusters albeit to modest levels (5, 7, 8 for IgG and 14 for IgM) (Fig. 3F, Fig. S7A, Table S6). Although TACI has been described in tonsil and BM ASC (*58*), we found high expression of TACI was seen in c14, a late IgM predominant ASC cluster (Table S6). In mice TACI expression is highest on mature innate-like B cells such as marginal zone and B-1 B cells which is critical for T-independent responses (*59, 60*). Healthy BM ASC do not express APRIL or BAFF showing the need for exogenous sources of these survival cytokines.

Other receptors in the TNFRSF considered to play roles in T cell activation were differentially regulated in early vs. late BM ASC clusters (Fig. 3F). For example, OX40 (*TNFRSF4*) and GITR (*TNFRSF18*) were both significantly upregulated in the late ASC clusters. In contrast, HVEM (*TNFRSF14*) expression was substantially increased in the early clusters 2, 3, 4, 9 and 13 (Fig. S7A, Table S6). In all, the sequential TNFRSF programs in early and late ASC provide important insights into BM maturation.

Although protein expression may be concordant with gene expression as shown with CD19 and CD138, not all surface markers may correlate with gene expression (*61*). By multi-dimensional flow analysis as BM ASC mature (downregulation of CD19 and upregulation of CD138), BCMA and TACI surface expression may be more variable and less concordant with gene expression (Fig. 4A). Based on our previous traditional BM subsets, pop A, B, and D (CD19^-^CD38^+^CD138^+^), both BCMA and TACI decrease surface protein expression with maturation in pop D. In contrast, as BM ASC mature, flow phenotyping revealed higher surface expression of OX40 and GITR based on the single cell data validating these expression profiles (Fig. 3F and Fig. 4B). Similarly, using the traditional subsets pop A, B, and D, OX40 and GITR increase in expression.

**Figure 4.**
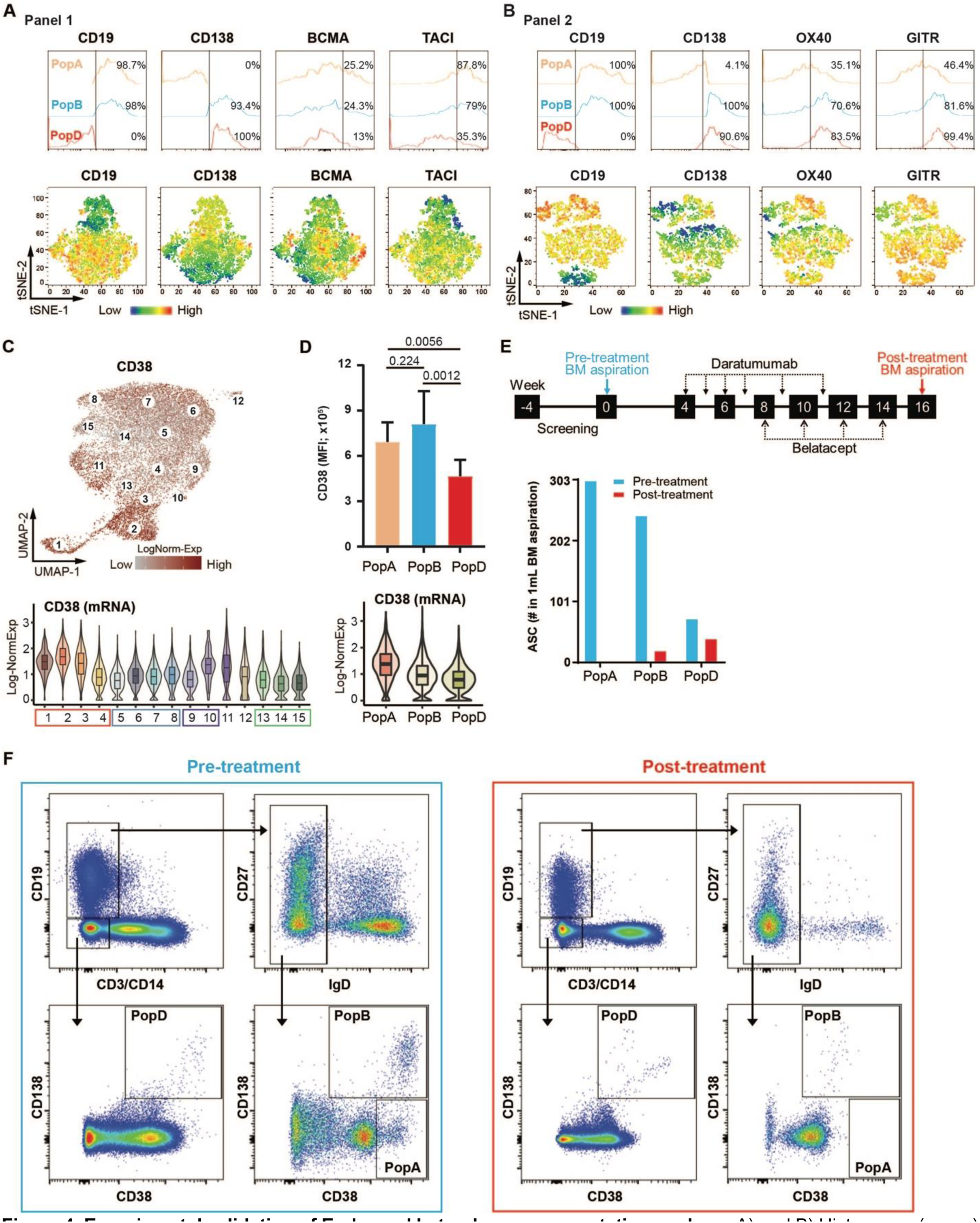
Experimental validation of Early- and Late-phase representative markers. A) and B) Histograms (upper) and tSNE heat maps (bottom) of expression of ASC surface markers (A) CD19, CD138, BCMA, and TACI (Panel 1), or (B) CD19, CD138, OX40, and GITR (Panel 2). C) CD38 log-Normalized gene expression visualized by UMAP (top) and violin plot grouped by cluster id (bottom). D) CD38 Mean Fluorescence Intensity (MFI) measured from n=9 healthy BM aspirates (top) and log-Normalized gene expression visualized violin plot grouped by FACS-sorted cell labels (bottom). E) Study regimen and timeline of the DSA patient (ClinicalTrials.gov Identifier: NCT04827979, top). Quantitation of the patient’s BM ASC subsets (# ASC per 1mL BM aspirate sample) of pre- and post-treatment with Daratumumab and Belatacept (bottom). F) Population gating of the patient’s BM sample. BM MNC were first gated for lymphocytes, singlets, and viability (based on their FSC/SSC and Live/Death properties). CD3 and CD14 were then used as dump markers to capture CD19+ and CD19-B cell populations. Subsequent sub-gating on the IgD-fraction (of CD19+ population) and using CD138 versus CD38 allow for breaking down BM ASC populations into 3 subsets of interest: PopA (CD19+CD38hiCD138-), PopB (CD19+CD38hiCD138+), and PopD (CD19-CD38hiCD138-).

Although CD38 expression increases greatly during B cell differentiation to ASC precursors (*62*), its downregulation during ASC maturation was unanticipated (Fig. 4C). Yet, this feature may actually be consistent with enhanced overall survival as shown in tumor cells (*63*). Similarly, pop D had lower MFI of CD38 surface expression by flow cytometry which was also concordant with gene expression of pop D cell (Fig.4D). Combined, these data may help understand the therapeutic targeting achieved by anti-CD38 agents currently used for the treatment of plasma cell malignancies and under investigation for highly sensitized patients and autoimmune conditions (*64–67*). Since we observed a downregulation of CD38 expression from early to late BM ASC clusters, we predicted a selective loss of early BM subsets with anti-CD38 therapy (Daratumumab). In a sensitized patient with a broad number of HLA antibodies waiting for 2^nd^ kidney transplant, we evaluated BM subsets pre- and post-treatment of a desensitization regimen of Daratumumab and Belatacept for 14 weeks (under ITN protocol ITN090ST, Fig. 4E). Of the 59 HLA-specific serum antibodies pre-treatment, only 16 remained post-treatment suggesting depletion of HLA-specific BM plasma cells. Using multi-color flow cytometry, we found dramatic depletion of immature BM ASC subsets, pop A and B with 100% and 92.2% reduction respectively (Fig. 4E,F). Only the late mature ASC subsets remained after treatment suggesting early phenotypes of BM ASC are the most susceptible to anti-CD38 therapies as predicted from the single cell analysis. Furthermore, a majority of HLA specificities may reside in the early compartments. In all, this unique single cell BM ASC analysis provides the first atlas and deep insights of BM plasma cell heterogeneity with implications for targeted therapeutic approaches.

#### Mature BM ASC downregulate pro-apoptotic genes and upregulate pro-survival genes

Previously reported bulk RNA-seq and ATAC-seq indicated that popD significantly up-regulates pro-survival genes *BCL2* and *MCL1*, despite enhanced chromatin accessibility being present only for *BCL2* (*6*). Additionally, popD showed significantly decreased expression of the pro-apoptotic genes *HRK*, *CASP3, BAX, BAK1* and *CASP8.* Our current data provide important replication of these features at the single cell RNA-seq level (Fig. S7C).

Extending this analysis, we examined the single cell expression of pro-survival, intrinsic and extrinsic pro-apoptotic genes, cell cycle, and cell cycle arrest across the 15 clusters (Fig. 5A). *TSC22D3*, *MCL1* and *BCL2* are the essential pro-survival genes for BM ASC maturation. *TSC22D3* (glucocorticoid-induced leucine zipper, GILZ) can inhibit the transcriptional activity of *FOXO3*, which leads to the further suppression of BIM-induced apoptosis, albeit in T cells (*68*). Of the late clusters, *TSC22D3* expression is high in c5 and c7 and the highest in c12 which has the corresponding highest expression of BIM (*BCL2L11*). *BCL2* is elevated to a similar degree in all late stage LLPC (c5-8), whereas *MCL1* shows even higher expression late in path2 (c7-8, Fig. 5A,B). Conversely, both intrinsic and extrinsic pro-apoptosis genes, including *BAX, BAK1, VDAC1, CASP3, CASP7* are reduced in the LLPC, again to similar extent in path1 and 2. These trends are consistent with a molecular basis for refractoriness to apoptosis in the late ASC (*69*).

**Figure 5.**
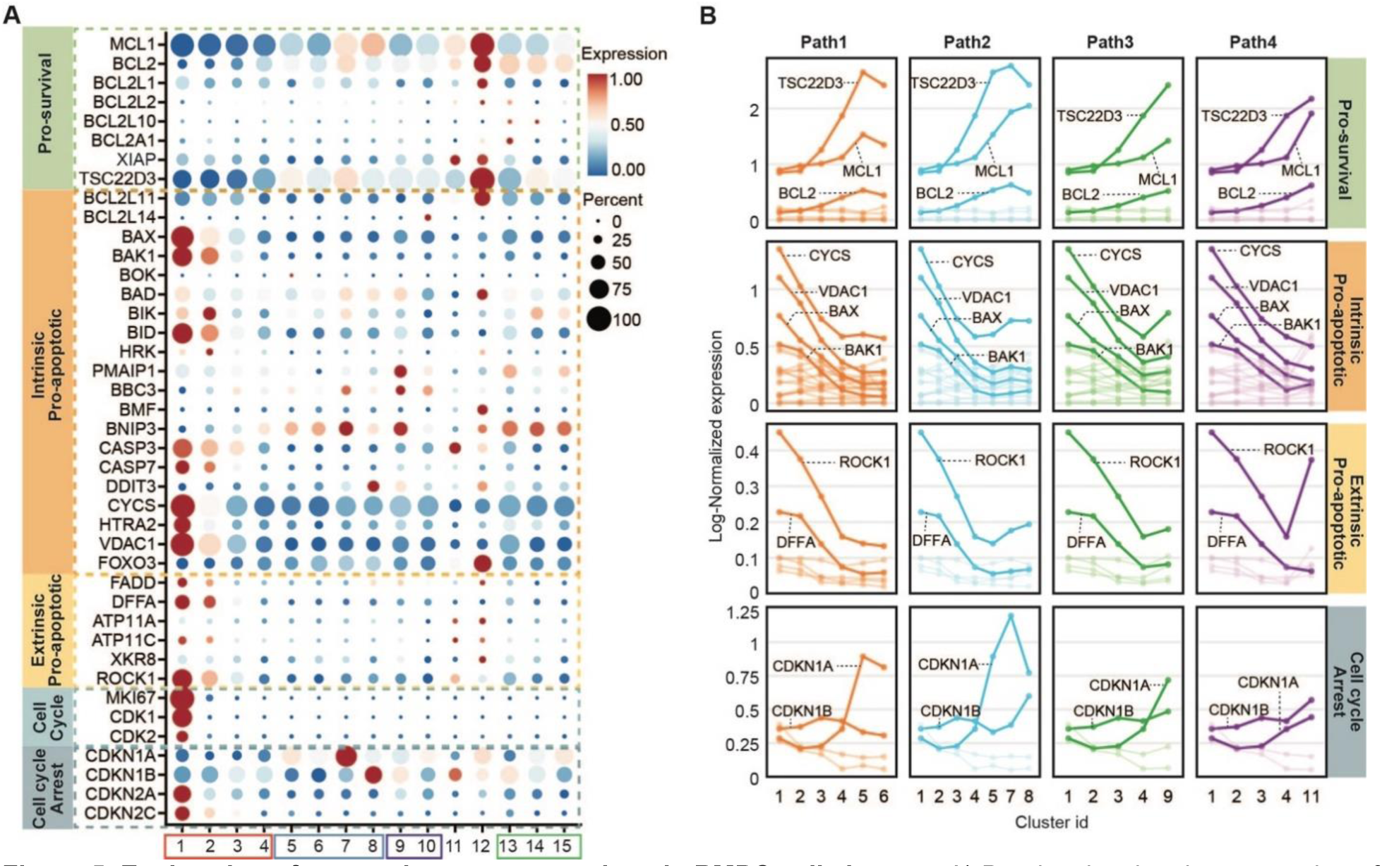
Exploration of apoptotic gene expressions in BMPC cell clusters. A) Dotplot showing the expression of genes related to pro-survival, intrinsic pro-, extrinsic pro-apoptotic, cell cycle progression and cell cycle arrest functions in BMPC subgroups. B) Dynamic gene expression alterations in 4 maturation paths defined in Fig.2B with the same color coding and ordering. The labeled thick solid lines showed the genes with high expressions and variations during the BMPC maturation.

Inability to mature from early to late ASC likely leads to cell death (Fig. 5A, B). Cluster 1 (PB) and cluster 2 stages of BM ASC maturation appeared to be primed for apoptosis with higher expression of pro-apoptotic genes, including the mitochondrial outer membrane permeabilization (MOMP) activators *BAX* and *BAK1* and mediator VDAC1; released apoptogenic proteins from the intermembrane of mitochondria cytochrome c (CYCS) and a serine protease OMI (encoded by *HTRA2*) that neutralizes the caspase-inhibitory proteins to directly and indirectly involve caspase activation. The DNA fragmentation factor (DFFA) and apoptotic membrane blebbing gene *ROCK1* were also upregulated (*70–73*). The PB stage also had high abundance of transcripts important for proliferation such as *MKI67*, *CDK1, CDK2* as well as cell cycle arrest associated genes p16 (*CDKN2A*) and p18 (*CDKN2C*). Two other cell cycle-inhibiting genes, p21 (*CDKN1A*), and p27 (*CDKN1B*) showed higher expression in clusters 7 and 8, respectively (*74*). Cell cycle arrest is largely regulated by activation of either one or both of the p16/pRB and p21/p53 pathways.

Cluster 6 showed high expression of *CD138* (SDC1), a member of the heparan-sulfate proteoglycan family which has been reported to promote ASC survival by regulating *BCL2* and *MCL1* (*75*). We also observed that *CD63*, *CD9* and *IL5RA* were up-regulated in the late stage BM ASC (Fig. S7D). *CD9* and *CD63* have been reported to associate with the metastatic ability of tumor cells, such that higher expression promotes decreased cell motility (*76*). IL6 is known to be important for BM ASC survival (*7, 77*), but interestingly IL6R expression was highest in the PB stage and cluster 2, whereas IL6ST (binding of IL6 and IL6R) showed higher gene expression in cluster 11 and 12, but not in the other late ASC clusters (Fig. S7D).

### Isotype characteristics of BM ASC

Our single cell analysis provided a thorough representation of the whole V(D)J repertoire of human BM ASC. Indeed, of the 17,347 cells with single cell transcriptomic information, 83% also had matched VDJ:V_H_ (by scVDJ-seq) sequences including isotype identification (Fig. 6A and Fig. S8A). The V_H_ repertoire included 11,853 clonal lineages, with a majority representing singletons (10,344 cells) while 1,509 lineages contained at least 2 cells (Fig. S8A). Of these, 421 were observed in only one of the 15 clusters, whereas 1,088 lineages were present in at least 2 cell clusters, mostly in adjacent UMAP spatial clusters (Fig. S8B). Each cell cluster contained a similar percentage of singletons except cluster 12 which only contributed a small number of cells and the highest proportion of non-matched VDJ cells (Figs. S8C, D). The polyclonal nature of the repertoire was not unexpected since BM ASC are the combined result of a lifetime of antigen exposures and reflect the historical serum antibody record.

**Figure 6.**
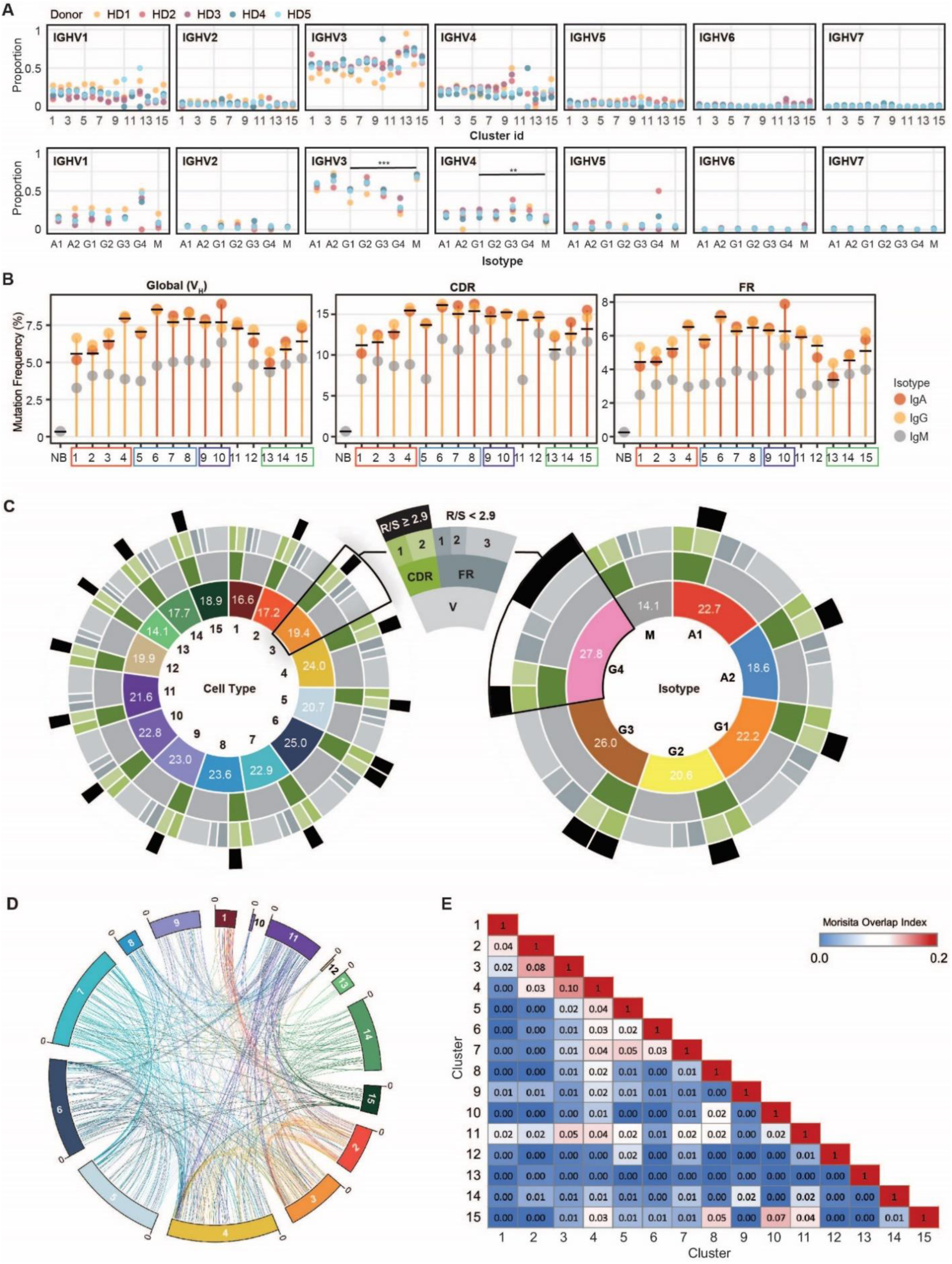
Mutation rate, similarity and connectivity of clones measured by scVDJ-seq. A) Summary of Ig heavy chain family gene usage. The y-axis shows the proportion of cells from each cell cluster (up) and isotype (down) have selected corresponding Ig heavy chain variable genes. Black bar showing the comparison objects and asterisks indicates the significance of statistical test (*** p ≤ 0.001, ** p ≤ 0.01). B) The average mutation frequency in the whole region V(Global), CDR and FR of IgA, IgG and IgM isotypes from each cell population. The solid black bar in each cell cluster indicates the overall average mutation frequency. NB is naive b cell extracting from one healthy donor’s peripheral blood and used as control. C) Summary of the average mutation numbers by scRNA-seq identified cell populations (left) and scVDJ-seq identified isotypes (right). The inner most circle shows the labels of the corresponding cell population or isotype. From inside-out, the colors of the first circle in each panel are consistent with cell colors defined in Fig. 1C) and 1G). Numbers in each section show the average number of mutations in region V. The second circle shows the average mutation numbers in CDR and FR of region V. The third circle further breaks CDR and FR into CDR1, CDR2, FR1, FR2, and FR3. The outer most circle, black bar indicates the selected region has replacement-to-silence (R/S) ratio greater than 2.9. D) Compiled Circos plot connecting individual subjects’ clones from cell clusters 1 to 15. Lineages were colored by the latest cell cluster. E) Heatmap showing the average Morisita overlap index of 5 subjects with 0 indicates no similarity and 1 indicates identical repertories. The redder the color indicates the higher similarity of clones between associated two cell clusters.

### Abundance of VH Ig transcripts per cell

Nearly every cell (98.6% ± 0.8%) had a consensus Ig gene with greater than 100 transcripts (Fig. S9). Single ASC in the transitional and late stage had a higher proportion of cells with large numbers of Ig transcripts (Fig. S9).

### V(D)J repertoire characteristics of BM ASC clusters

V_H_ gene usage was not biased between different ASC clusters, IgG isotypes, and individuals. Higher IGHV1, IGHV3 and IGHV4 family genes reflect the predicted distribution of these larger families (Fig. 6A). Similarly, there was no consistent bias of individual VH genes within the VH families.

The frequency and distribution of somatic hypermutation (SHM), provides important information regarding the maturation and antigenic selection of ASC and by extension, may offer important insight into their differentiation from separate B cell sources. As expected, BM ASC had higher global mutation frequencies relative to largely unmutated peripheral blood naive B (NB) cells from one healthy individual (Fig. 6B, NB = 0.33%, BM ASC = 6.89 ± 1.11%, p value _one-sided t test_ = 8.512e^-13^). Moreover, BM ASC displayed SHM frequencies comparable to historical values obtained from healthy human peripheral memory B cells (7.1%)(*78*). Interestingly, our BM ASC displayed SHM frequencies similar to those observed in blood ASC from vaccinated healthy control subjects (7.33%) but much higher than ASCs from patients with active autoimmune disease, systemic lupus erythematosus SLE (4.98%), a feature that has been ascribed to the large influx of newly generated naive-derived responses in the latter condition (*78*).

Overall, SHM frequencies ranged from 4.8% to 8.6% with the lowest frequency observed in c13 and the highest in c6 (Fig. 6B). IgG and IgA generally had significantly higher mutation frequencies than IgM isotypes but there was not a difference between IgA and IgG isotypes or IgG subclasses within each identified cell cluster (Fig. 6B; Table S7). Interestingly, the highest SHM mutation number was found in the small IgG4 fraction across clusters at 27.8 in average (Fig. 6C). Early-stage ASC of the IgG lineage, especially those with MHC class II gene expression (clusters 1-3) as well as all IgM lineage clusters, had a lower mutation frequency than late-stage IgG ASC. This result was evident across the global V region or within the framework region (FR) and complementarity-determining region (CDR), whether considered in isolation or combined (Fig. 6B). There was a trend toward higher average mutation rates as BM ASC mature suggesting either a potential survival advantage of cells with higher mutation frequencies or alternatively, preferential origination from late germinal center reactions. Cluster 6, which had the highest average number of mutations, also had the highest global V_H_, CDR and FR region mutation frequencies (permutation test, p < 0.05, Fig. S10).

Analysis of the regional distribution of SHM across the length of the VH gene within the framework region (FR) and complementarity-determining region (CDR), was also highly informative. Reflecting the length of the corresponding segment, average mutation numbers were higher in the FR than CDR, especially for FR3. A replacement-silent (R/S) mutation ratio greater than 2.9 is suggestive of antigen-selection (*79, 80*) and most of the clusters (14/15) had a CDR2 R/S >2.9. Only cluster 6 showed a R/S>2.9 in both CDR1 and CDR2, while cluster 1 had no regions with R/S > 2.9 (Fig. 6C). Overall, VDJ analysis showed that most BM ASC are highly mutated and antigen-selected.

#### Repertoire connectivity between early and late stage BM ASC

Among the 5 individual, we identified 11,853 V_H_ lineages, and all lineages were shared within a single individual demonstrating no public clonotypes. Clonal lineages were defined as the same V, J, CDR3 length, and 85% CDR3 homology. Of the 11,853 V_H_ lineages identified, 1,088 (9.2%), were present in at least 2 ASC clusters, thereby indicating a significant level of cluster interrelationship (Fig. S11B). However, to track the identical clone we used 98% homology to follow the same “clone” in the pseudotrajectories. Since IgG1ASC are the dominant isotype in the BM, we show the connectivity of identical clones of IgG1 ASC in clusters 1-15 in aggregate and by individual subject (Fig. 6D, Fig. S11A).

We applied the Morisita overlap index to map lineage connectivity across populations. Cell populations were highly connected within the pre-late (clusters 1-4) or the late stage (cluster 5-8) (Figs. 6D,E). The degree of connectivity was substantially greater after removal of singletons, a strategy that enriches for larger clones and increases the sensitivity of detection that otherwise would be necessarily limited by the number of cells available for analysis (Figs. S11B, C). Circos plots of IgG1 dominant cluster 1-8 are shown together with all clusters and all isotypes at 98% homology (Fig. S12). Using 98% homology to define identical clones, we found ASC in multiple cluster trajectories with Cluster 4 displaying the high connectivity with both early and late stage cell clusters further validating the inference that it is a “transitional” cluster by independent V_H_ repertoire analysis (Fig. S11). Interestingly, clusters 9 and 11 also showed connectivity with both early and late clusters (Fig. S11). Similarly, IgM predominant cluster 14 show high connectivity between corresponding IgM clusters 13 and 15 (Fig. 6E, Fig. S11), which together with single cell trajectories, denoted an evolutionary relationshhip between these subsets.

Although some identical clones were detected in multiple cell clusters, one of the biggest lineages 334 contained 11 cells distributed across early and late phase of IgG ASC as well as cluster 9,10, 11 and 15 (Fig. 7A). The VDJ sequences in this clone were identical as shown in the alignment (Fig 7B). There were several clones with 8 cells with identical sequences (385 and 280) that progressed through the early to late phases, validating potential paths for ASC maturation (Fig. 7A, B). While these observations show representative clones, many other inter-cluster clones contained identical sequences as well. These observations are consistent with a dynamic BM ASC compartment comprised of clones that likely undergo further maturation in the BM microniche.

**Figure 7.**
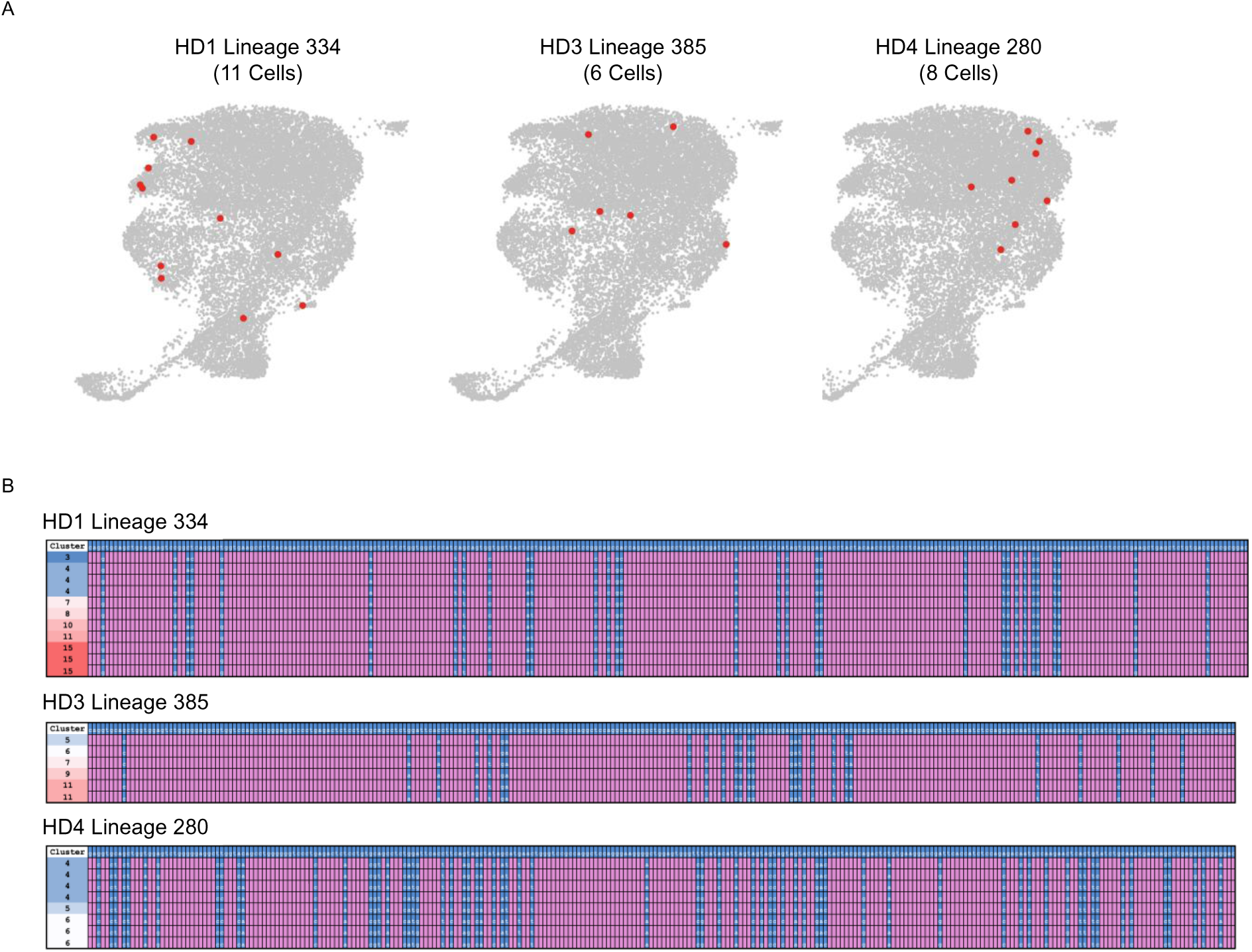
Cellular representation and alignment of clonal VH sequences that span multiple clusters. A) Clones were clustered by identical V, J, CDR3 length and 98% CDR3 homology to identify near identical clones and plotted on UMAP. Cells highlighted in red are members of the same clone. B) V region genes from the same clones were aligned against germline sequences. Blue squares indicate mutations in the individual sequences when compared to germline sequences and pink squares indicate matching nucleotides to the germline sequence. The cluster of each cell where the transcript was identified to be from is shown on the left.

## DISCUSSION

Antibody-secreting cells (ASC), represent a highly specialized effector compartment whose main physiological function is the production of protective antibodies, which in some cases, may persist for the lifespan of the host, owing to the existence of long-lived plasma cells (LLPC). While active infections trigger new antibody production through newly made ASC, either from naive or pre-existing memory B cells, long-standing antibodies represent a person’s immune history and contribute serological memory capable of preventing or moderating infection by previous agents. In contrast to other immune effector cells that perform their function during acute activation and are short-lived, LLPC are unique in that they constitutively perform their effector function while in a resting state and are endowed with prolonged survival. In contrast, short-term ASC, frequently referred to as plasmablasts, perform the same effector function while proliferating and may follow two different fates: post-proliferative apoptosis or subsequent differentiation into LLPC. It is well established that LLPC persist in specialized survival microniches in the bone marrow (BM). Yet, it has remained uncertain whether they arrive in the BM as fully differentiated cells that only require admission to the protective niche or instead, begin their BM journey as immature plasma cells that undergo additional differentiation. In the latter scenario, it remains to be determined whether all new ASC arrivals to the BM represent potential LLPC precursors, or alternatively, LLPC derive only from specific ASC subsets. In either situation, critical outstanding questions include: the specific phenotype of LLPC among mature BM ASC; the differentiation trajectories of BM ASC into LLPC; and their regulatory and survival programs.

While LLPC identity is commonly adjudicated by BM location (*81*), a significant degree of heterogeneity of ASC in that locale, would argue against that notion. Indeed, we previously defined 4 major BM ASC populations (A to D), on the basis of the expression of CD19 and CD138 and determined that LLPC with the ability to sustain anti-viral antibodies for over 4 decades, exclusively reside in CD19-CD138+ ASC or Pop D (*3*). However, the contribution of other ASC populations to long-lived response of lesser longevity (under 10 years of half-life for the corresponding serum antibodies such as anti-tetanus or anti-influenza), and the developmental relationships between different populations have remained unknown. Also undetermined has been the heterogeneity of the more mature pop D, as simply the acquisition of the corresponding surface phenotype may not necessarily correlate with prolonged longevity.

Our single cell studies provide an in-depth analysis of human BM ASC, thereby contributing substantial original insight into these questions. We provide the first high-resolution atlas of this immune compartment that define a high degree of BM ASC heterogeneity. For example, these studies offer an in depth understanding of the heterogeneity of BM ASC subsets which have tremendous implications of known (i.e anti-CD38 targets) and discovery of novel therapeutic mature BM plasma cell targets such as GITR and OX40. Moreover, our studies uncover maturation trajectories and delineate a developmental, regulatory, and survival roadmap of how to become a long-lived plasma cell that highlights interactions within the BM microniche.

From a developmental standpoint, we identified 15 distinct subsets that span early (c1-3) and late ASC populations (c5-8) with a small intermediate population (c4). The combination of KI67 and MHC class II expression distinguished early and late ASC with HLA-DOB and HLA-C providing resolution of late ASC beyond the broader CD19^-^CD38^+^CD138^+^ phenotype.

A finding of central significance is the identification of IgM bone marrow ASC clusters with properties of either early ASC (c13) or late ASC (c14-15). While long-term IgM memory B cells are well established as a first layer of memory responses upon which isotype-switched memory and ASC responses of higher affinity can be built (*82*), the ability to generate mature IgM LLPC has not been explored. To our knowledge, our results provide the definitive identification of mature BM PC of an IgM isotype, a population previously described in the mouse spleen as playing an important role on persistent immunity (*30*). Combined, trajectory studies and a substantial degree of clonal connectivity across these clusters, demonstrate not only the presence of mature IgM BM PC that might contribute long-lasting serum IgM antibodies but also suggest a process of local progressive differentiation of IgM and IgG ASC within the BM as they mature from early arrivals into terminally differentiated PC. Our data suggest that such differentiation may be regulated by similar maturation and survival programs acting on ASC generated from separate precursors. Future studies will be required to determine the precise contribution of extra-follicular (EF) and germinal-center (GC) driven pathways to BM IgM PC. An important feature of the IgM BM ASC is the presence of a substantial degree of SHM, a feature typically ascribed to GC-derived cells, yet also possible to generate in EF reactions as shown abundantly in the mouse and by our own work in humans (*30, 83–85*).

Once they arrive in the BM, ASC follow two major differentiation trajectories leading to the more mature BM PC populations. These two paths were distinguished by differences of TNF signaling through NFκB, which separates the late BM ASC trajectories. AP-1 factors are also of particular note among differentially upregulated genes in path2 (*JUN*, *JUNB*, *JUND*, *FOS*, *FOSB*, *MCL1*, *ZFP36L2*, *ELL2*, *CDKN1A*, *PRDM1*, *NFKBIA*, and others). Thus, JUNB, is involved in proliferation and survival(*86*), in diseased plasma cells (multiple myeloma, MM), but is abundantly expressed in healthy ASC where it likely plays an anti-apoptotic role. Similarly, while c-JUN is important for caspase-mediated c-ABL cleavage inducing apoptosis in MM, it may function quite differently in healthy BM ASC. Whether the role for the NF-κB/AP-1/STAT3 inflammatory regulatory network described in the transformation human cancers (*87*) also plays a role in LLPC will require further study.

Trajectory analysis also points to an important role for cell cycle regulation in late progression to terminally differentiated PC through path 2. Thus, p21, a marker capable of inhibiting a range of CDKs, is activated in a p53-dependent manner and was upregulated in the late phase cell cluster 7 to mediate cell cycle arrest (*88*). In turn, the tumor repressor p16, has also been described as a hallmark of cellular senescence (*89–91*). Terminal differentiation of cells that exit the cell cycle irreversibly and convert into specialized effector cells (*92*) are unique features of cellular senescence and potential mechanisms of LLPC survival.

Also of interest, path 2 was characterized by higher expression of ZFP36L1, an RNA-binding protein (RBPs), recently shown to facilitate newly-formed ASCs homing to the BM microniche (*93*). Although a role of RBP in ASC differentiation is unclear, these proteins may actually modulate ASC migration and survival in late ASC.

Our study also provides insight into the metabolic and survival underpinnings of late-stage BM ASC. From a metabolic standpoint, as they enter cell cycle arrest, downregulation of oxidative phosphorylation and fatty acid metabolism undergird differentiation into LLPC, together with upregulation of TNFα signaling via NFκB and pro-survival pathways. Of particular interest for the latter property is the differential expression of receptors belonging to the TNFRSF family, which transduce essential external survival signals delivered by cytokines known to regulate B cell and PC survival. Specifically, BCMA, binds APRIL to enhance survival through MCL-1 in mice and to prolong survival in human LLPC *in vitro* BM mimetic systems (*53, 56*). Interestingly, while BCMA was expressed across all BM ASC, it experienced a two-fold decrease in the more mature ASC, thereby pointing to an important role in securing survival through early differentiation. In contrast, TACI transcription experienced a modest increase from early to late BM ASC, a finding consistent with its heterogeneous expression in MM (*94*), and indicative of an unrecognized role in physiological LLPC maintenance. Although TACI is important for mouse ASC survival (*95*), its actual role and degree of redundancy with BCMA remains to be understood in humans. Given that TACI binds APRIL and BAFF with similar affinity, the current use of BAFF and APRIL inhibitors in autoimmune and malignant diseases and the interest in BAFF and BCMA CAR-T cells (*96*), our data should be helpful to inform the design and targeting of therapeutic agents to the ASC of interest. Finally, additional TNFRSF members including *HVEM*, *OX40*, and *GITR* are differentially expressed. *HVEM* appears important in early ASC, suggesting a role for LIGHT or BTLA, ligands for HVEM, in early ASC with implications in proliferation and cell-cycle arrest. Finally, cell-cell contact through OX40 and GITR which are only expressed in late ASC intimates a model where LLPC may have limited motility due to cell-cell contact as suggested in recent mouse studies (*97*).

In our highly-sensitized patients treated with Daratumumab and Belatacept, Daratumumab likely resulted in the specific loss of early BM ASC subsets; however, we cannot totally rule out the role of Belatacept inhibiting newly generated ASC immigrants into the BM site. Despite the importance of new ASC into the BM sites, it likely contributes only a small proportion of BM ASC within a 16-week period. Clearly, additional studies with single agents will be needed.

Finally, our data describe a detailed landscape of the intrinsic pro-survival and pro-apoptotic balance characteristic of early and late human BM ASC. Progressive upregulation of pro-survival genes in all 4 trajectory paths suggest the possibility of enhanced longevity; however, only path 1, 2, and 3 have continued downregulation of both intrinsic and extrinsic pro-apoptotic genes. Whether there is also closure of the pro-apoptotic regions as previously noted will require additional studies.

A significant limitation of our analysis is that it is restricted to a single snapshot in time of the LLPC BM maturation trajectories. Longitudinal analysis and clonal tracking in the course of an immune response would provide conclusive characterization of the precise identity and molecular programs of LLPC. This exercise however may be unpractical owing to the need for repeated sampling of the human BM compartment over several years. However, kinetics of maturation trajectories within our *in vitro* BM mimetic culture (*6*), together with scRNA-seq analysis should offer a strong experimental approach for the validation of these maturation programs.

Our study offers major advances to the field of LLPC biology. First, to overcome a major hurdle in the study of human PC, we applied a new specialized media to optimize the isolation and viability of single cell ASC (*7*). Combined with a sort-based enrichment step, this approach enabled the study of large numbers of rare and fragile BM ASC, which typically represent a very low fraction of BM mononuclear cells (*98*), a barrier previously overcome only for the single cell study of abundant circulating post-immunization plasmablasts (*99*). Secondly, our experimental and analytical design achieved mRNA sequencing of sufficient depth for quantitative measurement of non-Ig genes, a goal readily compromised by the abundance of Ig transcripts representing up to 90% of some BM ASC total transcripts. Finally, we utilized novel analytical pipelines to integrate human ASC transcriptomes and VDJ repertoires and to delineate the molecular roadmap of healthy LLPC maturation in the BM microenvironment.

Combined, our findings demonstrate substantial heterogeneity of mature BM ASC and suggest that ASC of LLPC potential may be present in several BM compartments. One possible interpretation is that actual LLPC might be restricted to specific late clusters. This possibility needs to be formally tested by localization of antigen-specific ASC to the different clusters as previously done by our groups with the broader ASC populations. This goal could also be accomplished through the transcriptomes of identifying single ASC of predetermined specificity.

## MATERIALS AND METHODS

### Sample recruitment

Healthy adults (n = 5) were enrolled at Emory University between 2018 – 2019. The mean age was 25± 4 [SD] years old with all of them were female. All studies were approved by the Institutional Review Board at Emory University and informed consent was provided.

### Cell sorting and library construction

Bone marrow aspirate was obtained under sterile conditions from the iliac crest from each of the 5 healthy adults. Mononuclear cells were isolated by Ficoll density gradient centrifugation and enriched by a custom designed negative selection EasySep cell isolation kit from StemCell that removes CD66b+/GPA+/CD3+/CD14+ cells to limit sorting time of fragile BM ASC. Enriched mononuclear cells were stained with the following anti-human antibodies: IgD–FITC (Cat. no. BD555778; BD Biosciences), CD3-BV711 (Cat. no. 317328; BioLegend), CD14-BV711 (Cat. no. 301838; BioLegend), CD19-PE-Cy7 (Cat. no. 301838; BD Biosciences), CD38-V450 (Cat. no. BDB561378; BD Biosciences), CD138-APC (Cat. no. 130-117-395; Miltenyi Biotech), CD27-APC-e780 (Cat. no. 5016160; eBiosciences), and LiveDead (L34966; Invitrogen). ASC subsets were FACS-sorted on a BD FACSAria II using a standardized sorting procedure with rainbow calibration particles to ensure consistency of sorts between individuals. ASC subsets were sorted as: popA (CD19^+^CD38^hi^CD138^-^), popB (CD19^+^CD38^hi^CD138^+^) and popD (CD19^-^CD38^hi^CD138^-^). Up to 17,000 cells were FACS-sorted from each population into RPMI with 5% FBS to maintain viability.

FACS-sorted cells were kept on ice until proceeding with 10x Genomics processing. Cells were centrifuged at 500xg for 10 minutes at 4C to remove media. Cells were then washed twice in 0.04% BSA in PBS. During last wash, media was aspirated and cell volume was measured using a pipette. The whole sample was then taken for processing using the 10x Genomics 5’ v1 Single Cell platform. V(D)J and 5’ Gene Expression (GEX) libraries were constructed for each sample, following 10x Genomics protocol. QC for each library was performed at each step by Bioanalyzer. Final libraries were quantified by kappa qPCR and sequenced on a NovaSeq.

### Pre-processing of 10x Genomics scRNA-seq data and quality control

The 10x Genomics single-cell transcriptomic sequenced raw reads were aligned to GRCh38 reference and quantified per cell barcode using Cellranger v3.0.1 (https://support.10xgenomics.com/single-cell-gene-expression/software/downloads/latest). Genes expressed in at least 0.1% - 0.3% (depending on the sample size) of the total cell population were regarded as expressed genes and retained for downstream analysis. To filter low-quality cells, we excluded cells with ≥ 30% of their UMIs coming from mitochondrial genes, or ≤ 800 total number of detected genes, or total number of UMIs ≤ 1000. We first retained cells with a detected gene number between 200 – 800 (labeled as LowGN in Fig. S1A), but none of these genes were significantly expressed in the cell group. Moreover, we removed cells with total number of genes or UMIs ≥ 6,000 or 60,000, respectively, to control for potential doublets. Additionally, we filtered out cells with immunoglobulin genes corresponding UMI count percentage ≤ 5% to avoid contaminated non-antibody secreting cells. Then, we excluded contaminated cells expressing diagnostic cell markers, eg. CD3E (T cell), CD16 (encoded by *FCGR3A*) and CD14 (Monocytes), NKG7, GNLY (Natural killer cells), HBB (Erythrocytes), CD20 (encoded by *MS4A1*), PAX5, IRF8 (B cells).

### Normalization and cell cluster detection

The scRNA-seq data was next analyzed using version 3.2.2 of the Seurat package(*100, 101*). The gene expression counts of each cell were normalized using regularized negative binomial regression (SCTransform) to account for sequencing depth(*102*). Next, we selected the top 3,000 highly variable features (HVFs) based on their standardized expected variance after variance-stabilizing transformation, removed all the immunoglobulin genes from the HVF list, and then used HVF data from SCTransform residuals to perform principal component analysis (PCA). In the first run of our scRNA-seq analysis pipeline, we falsely included 2 misannotated Ig genes AC233755.1 and AC233755.2 in the HVF list, which resulted in 15 clusters plus 2 additional AC gene-driven cell clusters (Fig. S2B). Subsequently, we further excluded these two genes before cell clusters detection. Next, we utilized Canonical Correlation Analysis (CCA) from r package Seurat to anchor each dataset, removing individual effect and generating an integrated dataset (*100*). Using this CCA-integrated dataset, a graph-based clustering method was applied to build a shared nearest neighbor (SNN) graph in PCA space, after which the Louvain community identification algorithm was applied to group cells at the set resolutions, with higher values leading to a greater number of clusters (*103*).To assess the stability of the clustering based on different combinations of running parameters (dimension numbers = 50, 60 and 70, PC numbers = 30, 50, and 70 and resolution parameter = 0.2, 0.5, 1.0 and 1.5), we computed the Rand Index (RI) between pairs of classifications derived with different parameters. RI is a similarity measurement taking values from 0 (low) to 1 (high), which computes the proportion of cell pairs that are in agreement between cell clusters from two different parameters. Finally, we used 70 dimensions to anchor individual datasets and account for subject differences, and 50 PC were used to construct a SNN graph to detect cell clusters (*101, 104*). A resolution of 1.0 gave the most stable cell clusters (Fig. S2D). 15 cell clusters from the first run were retained with the cells from the two AC gene-driven clusters dispersed throughout the remaining 15 clusters. Finally, these clusters were visualized in two dimensions using uniform manifold approximation and projection (UMAP)(*105*).

### Differentially expressed and marker gene detection

Differentially expressed genes (DEG) between two cell clusters were identified based on LogNormalize SCTransform-adjusted count matrix, which is from data slot of SCTransform function in the Seurat r package. It’s worth noting that the HVF list with excluded Ig genes was used to detect cell clusters, the whole gene expression list which included Ig genes was used to detect cell markers and DEGs. It was performed by using zero-inflated generalized linear models including individual as a random covariate, with the MAST package (*106*), which was proven effectiveness on both real measured and simulated single-cell data (*107*). Cell cluster marker gene detection used the same settings but compared gene expression between cells from each selected cell cluster versus all remaining cells from the other clusters. Marker genes were defined as significantly up-regulated genes in associated cell clusters with average natural-log fold change (avgLogFC) greater than 0.25 and Bonferroni adjusted p value less than 0.05, while top DEGs were also included down-regulated genes with avgLogFC less than -0.25.

### GSEA Hallmark enrichment analysis

For gene set enrichment analysis, all expressed genes were ranked in descending order by multiplying the negative log-p value (NLP) derived from DEG analysis by the sign of the avgLogFC between the two clusters. The resultant pre-ranked gene list was used as input into GSEA v4.0.3 Preranked analysis (*108*). The enrichment score is derived from a multivariate U score (*109, 110*). Briefly, scores were calculated by averaging normalized expression levels for all the transcripts that were identified as maturation-associated DEGs, which were differentially expressed in any two adjacent stages of plasma cell maturation in Fig. 1C or differentially expressed between early and late cell clusters and annotated in the selected HALLMARK pathways.

### Trajectory analysis

For the IgG dominant cell trajectory analysis, we used the slingshot method as it can detect the bifurcation, multifurcation, linear and tree-like differentiation topology (*111*). We ran slingshot r package v1.4.0 (*111*) with UMAP embeddings as input data, cell cluster ids as cell labels and cluster 1 (PB) as the starting point, and all other values using default settings. Although numerous methods have been developed for single cell trajectory imputation, strikingly, out of the 45 methods reviewed by Saelens et al. (*112*), only three (PAGA and RaceID/StemID) can be used to detect disconnected graphs, but none of these releases are currently stable. We assumed that each lineage had its own progenitor population, so for the accuracy and stability of inference, we focused on dissecting the maturation paths only for IgG lineage cell populations which are likely to originate with the PB (cluster 1), whereas the progenitor for IgM remains unclear.

### Transcription factor analysis

Transcription factors were combined from AnimalTFDB (*113*) and known human transcription factors (*114*). For downstream analysis, we only focused on the TFs that were either identified as HVFs or markers of any one of cell clusters or differentially expressed in the comparison of any adjacent two clusters of PC maturation or between the early and late phases of PC maturation. Only those TFs expressed in at least 10% of any one of the BMPC subgroups were included in detection of cluster-specific TFs; and only those expressed in at least 20% of associated cell clusters and having avgLogFC greater than or equal to 0.25 between the two cell groups with the highest and lowest gene expression were selected for visualization. Averaging the expression of TFs by cluster id and defined maturation stage, each TF was assigned to the cell cluster or stage with the highest expression. For those TFs showing greater than or equal to 0.25 avgLogFC between assigned cell cluster and cell cluster with the second highest expression were labeled as cluster distinct. The same criterion was used to label stage-distinct TFs. Associations between pairs of TFs were evaluated using annotations in the STRING protein-protein interaction database (*40*); only interactions that have a combined score greater than 500 and exist within the assigned cell cluster were retained for visualization.

### Multicolor flow cytometry for experimental validation

MNC were isolated from 4 (Panel 1) and 2 (Panel 2) BM aspirate samples from heathy donors using a Ficoll density gradient and stained with the following anti-human antibodies: IgD-Brilliant Violet 480 (Cat. No. 566138; BD Biosciences), CD3-BUV737 (Cat. No. 612750; BD Biosciences), CD14-BUV737 (Cat. No. 612763; BD Biosciences), CD19-Spark NIR 685 (Cat. No. 302270; BioLegend), CD38-Brilliant Violet 785 (Cat. No. 303530; BioLegend), CD138-APC-R700 (Cat. No. 566050; BD Biosciences), CD27-Brilliant Violet 711 (Cat. No. 356430; BioLegend), BCMA-Brilliant Violet 421 (Cat. No. 357520; BioLegend), TACI-PE-Cy7 (Cat. No. 311908; BioLegend), OX40-Brilliant Violet 510 (Cat. No. 350026; BioLegend), GITR-Brilliant Violet 605 (Cat. No. 747664; BD Biosciences), and Zombie NIR Fixable Viability Kit (Cat. No. 423106; BioLegend). The anti-human antibodies that were used for staining BM MNC isolated from the patient with high donor-specific antibodies (DSA) awaiting kidney transplant in the ITN protocol (ClinicalTrials.gov Identifier: NCT04827979) include: IgD-BB700 (Cat. No. 566538; BD Biosciences), CD3-PE-Cy5 (Cat. No. 2363822; ThermoFisher), CD14-PE-Cy5 (Cat. No. 2319032; ThermoFisher), CD19-BUV395 (Cat. No. 740287; BD Biosciences), CD38-FITC (Cat. No. CYT-38F2-A; Cytognos), CD138-PE-Cy7 (Cat. No. 356514; BioLegend), CD27-BV605 (Cat. No. 562655; BD Biosciences), and Live/Dead (Cat. No. L34962; ThermoFisher). Cells were acquired on a Cytek Aurora spectral flow cytometer using Cytek SpectroFlo software (Cytek; the HD BM samples) or a LSR Fortessa X20 (special order research product with 5 lasers; BD Biosciences) and analyzed using FlowJo software (v10.8.1; with DownSample (v3.3.1) plugin; BD Biosciences).

### Single cell VDJ sequencing (scVDJ-seq) and analysis

Cells were counted using a Bio-Rad TC10 cell counter and verified via hemocytometer. Cell numbers were adjusted to 1,000 cells per μl to allow for 10,000 single cells per sample loaded in the 10x Genomics Chromium device. The manufacturer’s standard protocol for Chromium Next GEM Single Cell 5’ Reagent Kit, v2 and Chromium Single Cell Human BCR Amplification kit was used to generate libraries. Libraries were sequenced on an Illumina NovaSeq (paired-end; 2 × 150 bp; read 1:26 cycles; i7 index: 8 cycles, i5 index: 0 cycles; read 2: 98 cycles) such that more than 70% saturation could be achieved with a sequence depth of 5,000 reads per cell for VDJ libraries.

Analysis of single cell VDJ data was conducted using Cellranger v3.1.0 via the 10x Genomics cloud interface and an in-house developed informatics pipeline for clonal clustering and SHM analysis (*78*). Fasta files from the Cellranger output were annotated with metadata and aligned to germline sequences using the IMGT/HighV-QUEST web portal (*115*). All data from IMGT/HighV-QUEST were retained through the process and were used for mutation calculations and alignment analyses. The definition of lineages/clones is consistent with previous publication (*78*). The frequency and distribution of somatic hypermutation were ascertained on the basis of non-gap mismatches of expressed sequences with the closest germline V_H_ sequence. Mutation frequencies were determined by calculation of the number of mutations in V regions relative to the number of bases in non-gap V regions. The ratio of replacement mutations to silent mutations were calculated for CDR and framework regions separately from the corresponding V_H_ areas. In sequences with non-zero replacement but zero silent mutations, the number of silent mutations was set to 1 (*116*). Merging of gene expression and VDJ data sets and subsequent analysis was conducted using in-house developed pipelines in conjunction with the immcantation pipeline (*117*) and Seurat. Circular visualization plots were created with Circos software v0.69-6.

### Permutation test for mutation detection

In order to assess whether mutation numbers differ among clusters, we first randomly shuffled mutation frequency data in all the cells and grouped them using current cell clusters. We then performed ANOVA test and Tukey’s HSD (honest significant difference) consecutively. After repeating the previous steps one hundred times, we applied multiple comparison adjusted p values from the TukeyHSD test to calculate the p value for permutation test: P(permutation test) = (Number of permutation tests showing p value from TukeyHSD < p value obtained from running real data) /100.

## Supplementary Materials

Fig. S1. Integrated single-cell transcriptomic datasets.

Fig. S2. Stability and reliability of cell clusters.

Fig. S3. The features and pathways characterize the IgM dominant cell populations.

Fig. S4. Distribution of selected features in cell clusters.

Fig. S5. Patterns of Hallmark pathway enrichment analysis.

Fig. S6. Genes driven the difference of TNFα signaling via NFKB pathway in late LLPC.

Fig. S7. Visualization of indicative genes.

Fig. S8. Summary of scVDJ-seq data detected clones/lineages.

Fig. S9. Summary of Ig heavy chain gene transcript numbers in each cell cluster.

Fig. S10. Permutation results for the mutation frequency test.

Fig. S11. Clonal connectivity visualization in all cell clusters.

Fig. S12. Clonal connectivity of nearly identical clones in IgG1-dominant clusters 1 to 8.

Table S1. Cell cluster marker genes.

Table S2. Summarization of 205 differentially expressed TFs

Table S3. Comparison of top DEGs between 5vs6 and 5vs7.

Table S4. Comparison of GSEA preranked results between 5vs6 and 5vs7.

Table S5. HALLMARK_TNFA_SIGNALING_VIA_NFKB output results by running GSEA prerank analysis.

Table S6. TukeyHSD posthoc test for TNF family gene expression between compared two clusters.

Table S7. TukeyHSD posthoc test for the whole V (Global), CDR and FR regions mutation frequency among different isotypes.

## Acknowledgements

We thank Shuya Kyu of the Lee lab for technical assistance. We thank Robert E. Karaffa and Kametha T. Fife of the Emory University School of Medicine Flow Cytometry Core (EFCC) for technical support. We thank Kitza Williams for coordinating and managing the ITN project. We also thank our team of clinical coordinators and donors who made this study possible.

## Author contributions

Experimental design: FEL

scRNA-seq and scVDJ-seq data analysis: MD

scVDJ-seq data supervision: CMT, IS

Interpretation and visualization: MD

Validation experiments performance and analysis: DCN

Healthy human BM aspirates acquisition: SL, JA

Experiments performance: CS, CJJ, CK, IH, SK, SM

Experimental guidance: CJJ, DCN, AK, EG

ITN study supervision: RR, FV, AJ, SJK

ITN sample managing and processing: SC, TM, CB

Supervision: FEL, GG

Writing – original draft: MD, FEL

Writing – review & editing: MD, FEL, IS, GG

**Competing interests:** F.E.L. is the founder of Micro-Bplex, Inc., serves on the scientific board of Be Biopharma, is a recipient of grants from the BMGF and Genentech, Inc., and has served as a consultant for Astra Zeneca. I.S. has consulted for GSK, Pfizer, Kayverna, Johnson & Johnson, Celgene, Bristol Myer Squibb, and Visterra. F.E.L, DCN, & IS are inventors of the Issued patents: 9/21/21 US 11,124766 B2 PCT/US2016/036650 and 9/21/21 US 11, 125757 B2 for the plasma cell survival media. Provisional patents have been filed for identification of human LLPC biomarkers with inventors: FEL, MD, IS, GG. The other authors declare no competing interests.

