## Supplemental figures for "Human Bone Marrow Plasma Cell Atlas: Maturation and Survival Pathways Unraveled by Single Cell Analyses"

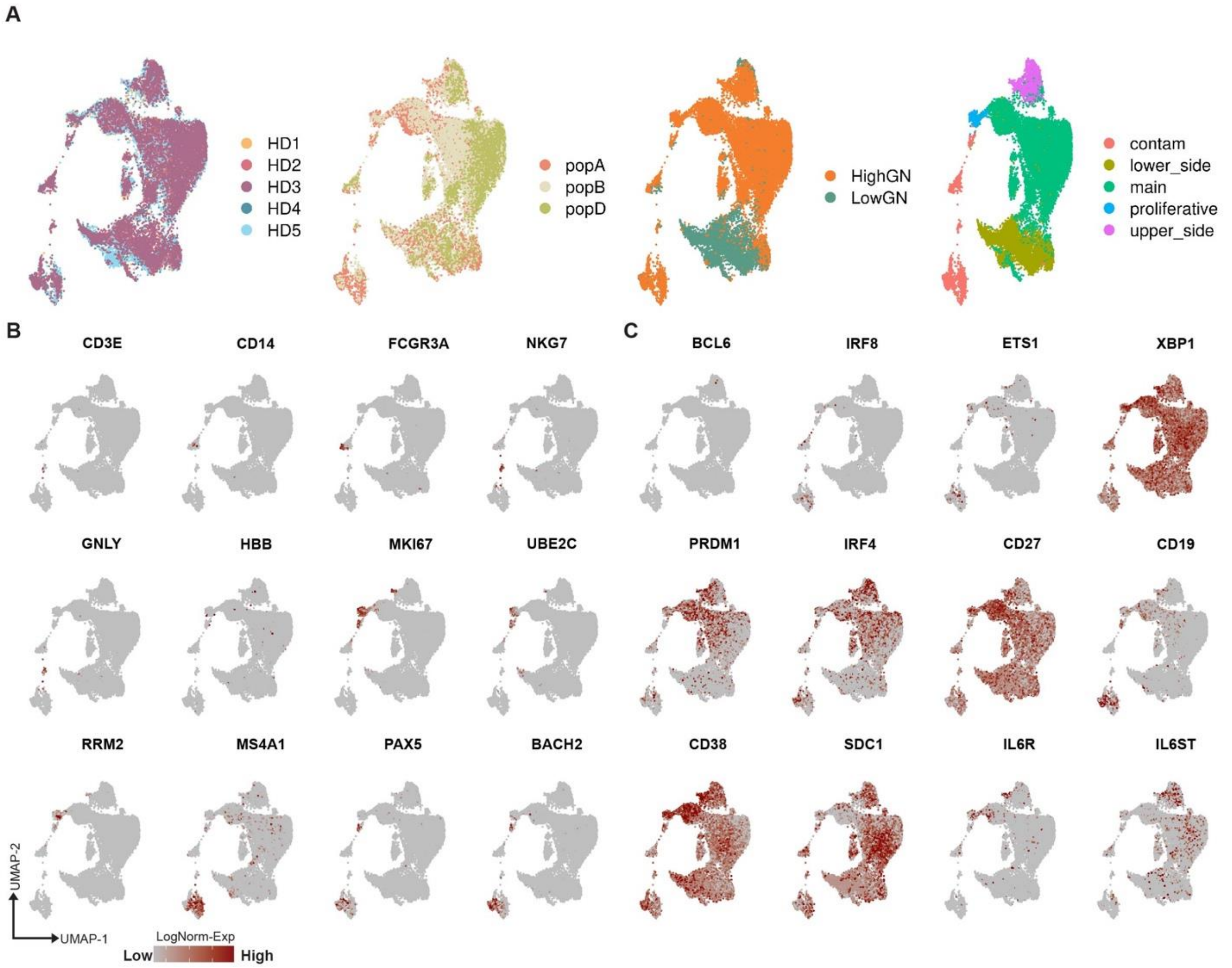


**Figure S1. Integrated single-cell transcriptomic datasets.** A) UMAP plots of single-cell transcriptomic profiles from 5 subjects before 2-step cell quality control and colored by sorted subject id, FACS-sorted labels, gene number group and cell subsets, which is mainly grouped by the location of cell population on UMAP, proliferating status and whether they are contaminating cells. HighGN: high gene number, indicating cells have greater than 800 detected genes, which is correspond to the gene cutoff in Fig. 1B; LowGN: low gene number, representing cells have greater than 200 detected genes but less or equal than 800 genes; contam: contaminating cell populations. B) and C) Contaminated cell population (B) and plasma cell (C) associated master gene expression. The redder the dot indicates the higher log-normalized gene expression.

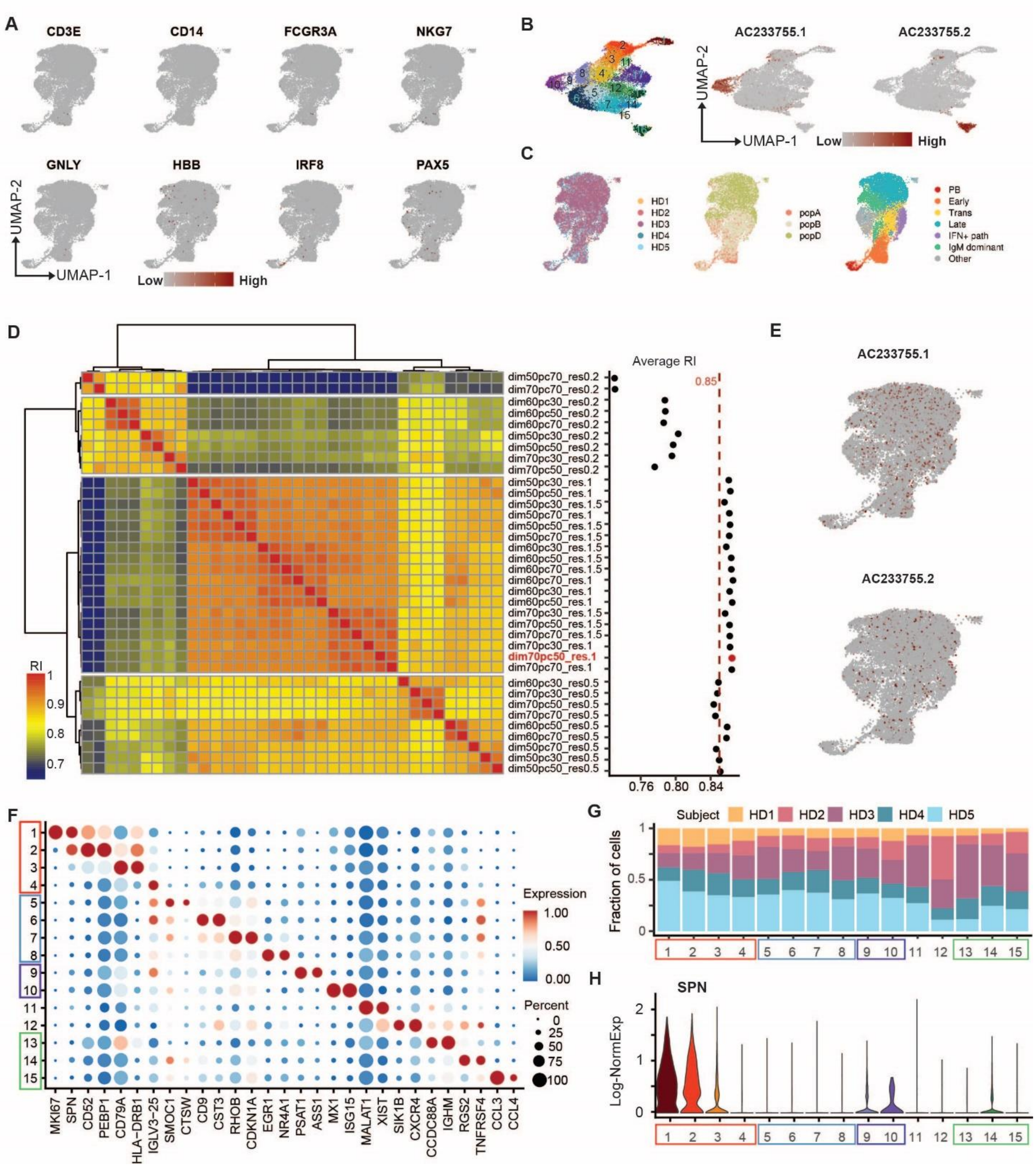


**Figure S2. Stability and reliability of cell clusters.** A) Expression of contaminated cell markers in current UMAP. B) Cell clusters (left) when not removing AC233755.1 and AC233755.2 genes before detecting cell clusters in the first run. Expression of AC genes in the previous UMAP (right). C) Current UMAP colored by sampled subjects, FACS-sorted cell populations and defined stages of BMPC maturation. D) Two-way hierarchical clustering of RI by comparing cell clustering results from running different combinations of key cell clustering parameters (left), the parameter information was shown in the middle labels and scatter plot showed the average RI of each row of heatmap (right). Dashed red line labeled the RI value 0.85. (dim: dimension; pc: PC; res: resolution.) E) Expression of AC genes in current UMAP. F) Dot plots for expression of marker genes in each subgroup of bone marrow plasma cell defined in Fig. 1C. Here and in later figures, colors represent Min-Max normalized mean expression of marker genes in each cluster, and dot sizes indicate the proportion of cells expressing each marker gene. G) Fraction of cells in current scRNA-seq data defined cell clusters by sampled subjects. H) Log-normalized gene expression of SPN in cell clusters.

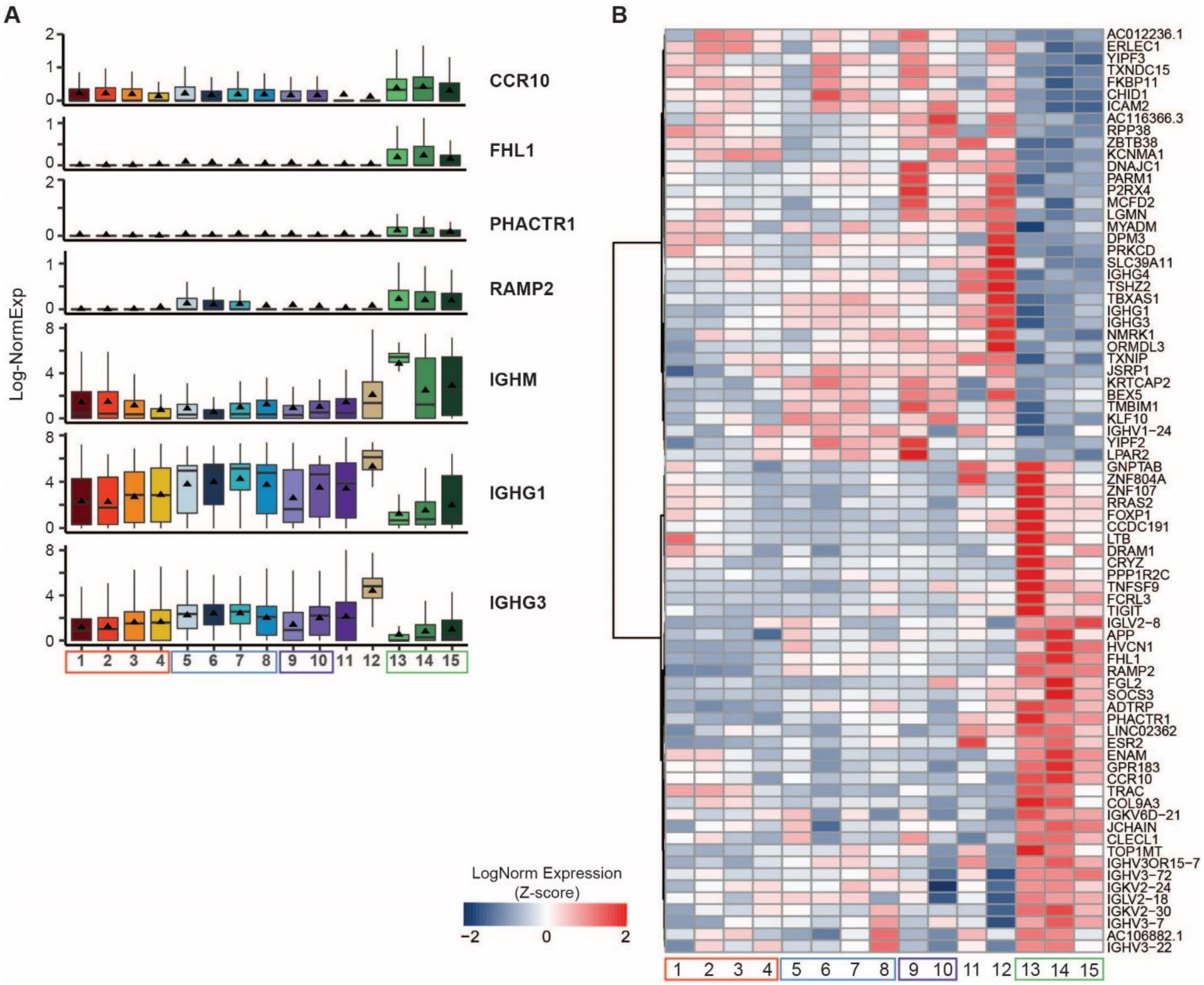


**Figure S3. The features and pathways characterize the IgM dominant cell populations.** A) Boxplots showing the expression of indicative IgM highly expressed genes as well as three Ig genes associated with somatic recombination of immune receptors in (C). B) Heatmap showed the row-scaled gene expression of CCR10 highly correlated genes (|Pearson correlation coefficient| ≥ 0.6, '||' implies absolute value).

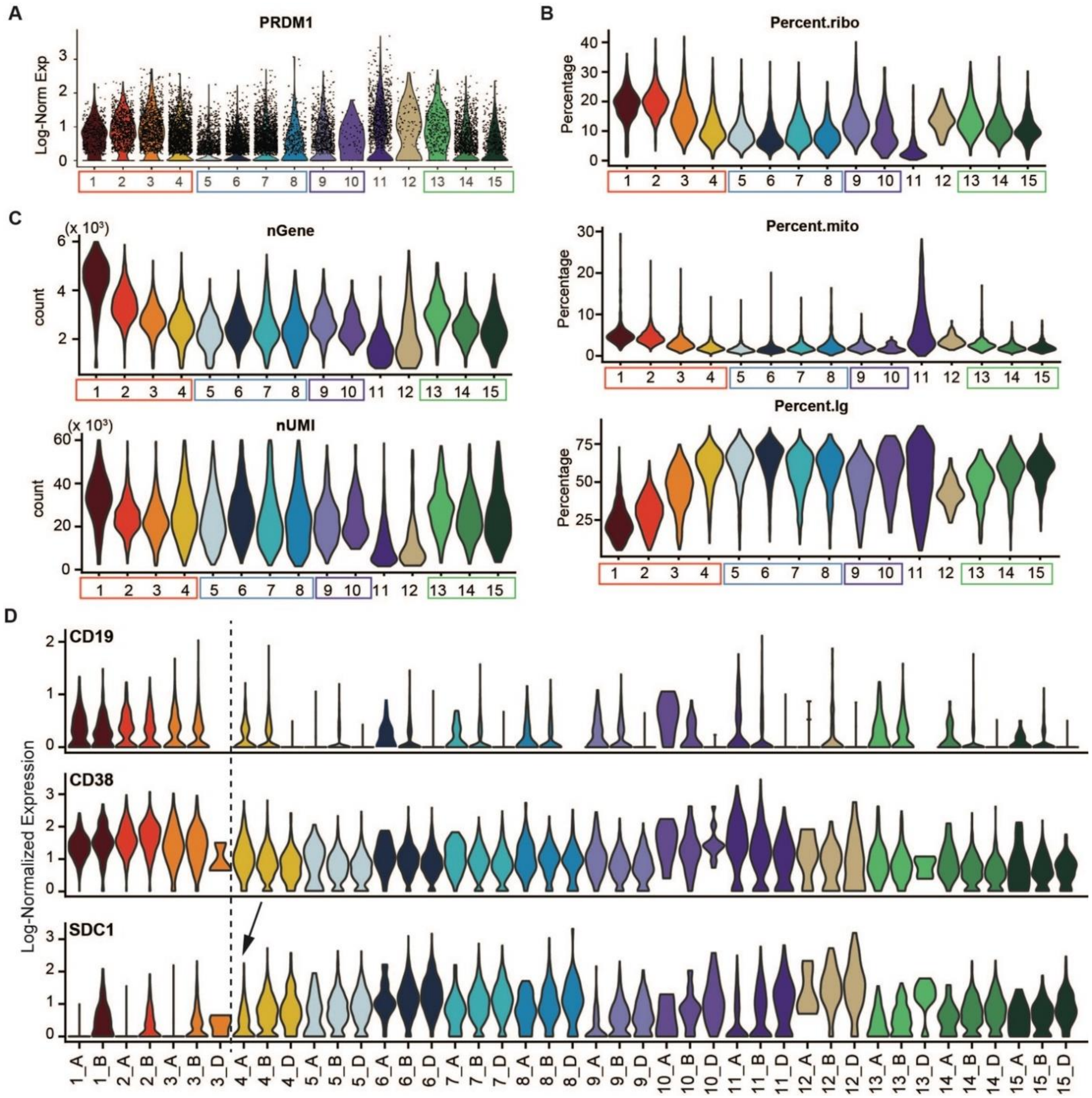


**Figure S4. Distribution of selected features in cell clusters.** A) Log-normalized gene expression of PRDM1 in cell clusters. B) and C) Violin plots show the percentage of transcripts coming from ribosomal genes (ribo), mitochondrial-encoded mitochondrial genes (mito), and immunoglobulin genes (Ig) (B) and the distribution of total detected gene numbers (nGene), total UMIs (nUMI) (C). D) X-axis is regrouped cell populations when taking both scRNA-seq defined cell subgroups and FACS-sorted cell labels into account. Y-axis is the log-normalized gene expression. Arrow points out the FACS-sorted popA cells starting CD138 (encoded by SDC1) transcripts.

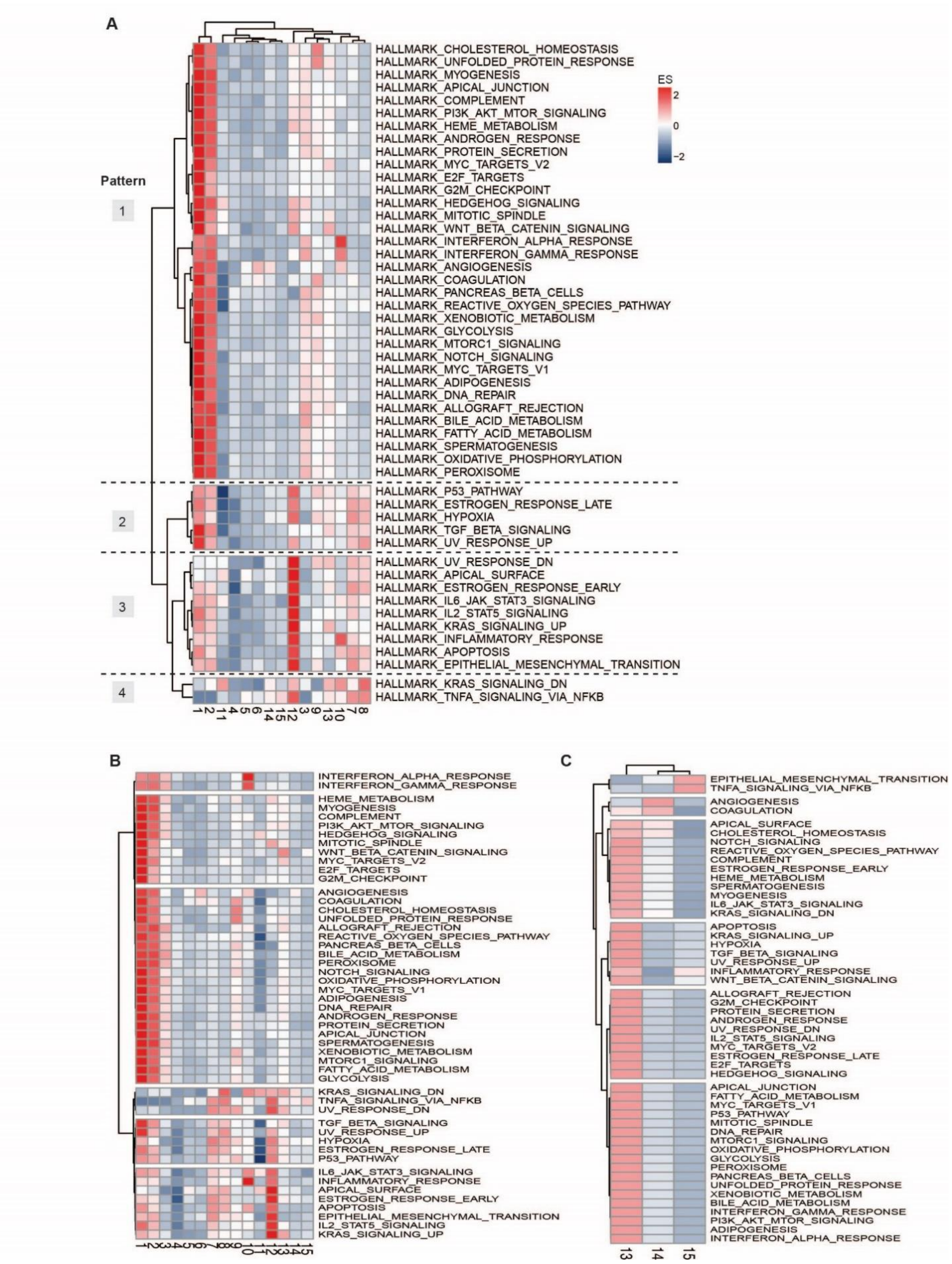
A
Pattern
1
2
3
4
ES
2
0
−2
HALLMARK_CHOLESTEROL_HOMEOSTASIS
HALLMARK_UNFOLDED_PROTEIN_RESPONSE
HALLMARK_MYOGENESIS
HALLMARK_APICAL_JUNCTION
HALLMARK_COMPLEMENT
HALLMARK_PI3K_AKT_MTOR_SIGNALING
HALLMARK_HEME_METABOLISM
HALLMARK_ANDROGEN_RESPONSE
HALLMARK_PROTEIN_SECRETION
HALLMARK_MYC_TARGETS_V2
HALLMARK_E2F_TARGETS
HALLMARK_G2M_CHECKPOINT
HALLMARK_HEDGEHOG_SIGNALING
HALLMARK_MITOTIC_SPINDLE
HALLMARK_WNT_BETA_CATENIN_SIGNALING
HALLMARK_INTERFERON_ALPHA_RESPONSE
HALLMARK_INTERFERON_GAMMA_RESPONSE
HALLMARK_ANGIOGENESIS
HALLMARK_COAGULATION
HALLMARK_PANCREAS_BETA_CELLS
HALLMARK_REACTIVE_OXYGEN_SPECIES_PATHWAY
HALLMARK_XENOBIOTIC_METABOLISM
HALLMARK_GLYCOLYSIS
HALLMARK_MTORC1_SIGNALING
HALLMARK_NOTCH_SIGNALING
HALLMARK_MYC_TARGETS_V1
HALLMARK_ADIPOGENESIS
HALLMARK_DNA_REPAIR
HALLMARK_ALLOGRAFT_REJECTION
HALLMARK_BILE_ACID_METABOLISM
HALLMARK_FATTY_ACID_METABOLISM
HALLMARK_SPERMATOGENESIS
HALLMARK_OXIDATIVE_PHOSPHORYLATION
HALLMARK_PEROXISOME
HALLMARK_P53_PATHWAY
HALLMARK_ESTROGEN_RESPONSE_LATE
HALLMARK_HYPOXIA
HALLMARK_TGF_BETA_SIGNALING
HALLMARK_UV_RESPONSE_UP
HALLMARK_UV_RESPONSE_DN
HALLMARK_APICAL_SURFACE
HALLMARK_ESTROGEN_RESPONSE_EARLY
HALLMARK_IL6_JAK_STAT3_SIGNALING
HALLMARK_IL2_STAT5_SIGNALING
HALLMARK_KRAS_SIGNALING_UP
HALLMARK_INFLAMMATORY_RESPONSE
HALLMARK_APOPTOSIS
HALLMARK_EPITHELIAL_MESENCHYMAL_TRANSITION
HALLMARK_KRAS_SIGNALING_DN
HALLMARK_TNFA_SIGNALING_VIA_NFKB
1 2 11 4 5 6 14 15 12 3 9 13 10 7 8
B
INTERFERON_ALPHA_RESPONSE
INTERFERON_GAMMA_RESPONSE
HEME_METABOLISM
MYOGENESIS
COMPLEMENT
PI3K_AKT_MTOR_SIGNALING
HEDGEHOG_SIGNALING
MITOTIC_SPINDLE
WNT_BETA_CATENIN_SIGNALING
MYC_TARGETS_V2
E2F_TARGETS
G2M_CHECKPOINT
ANGIOGENESIS
COAGULATION
CHOLESTEROL_HOMEOSTASIS
UNFOLDED_PROTEIN_RESPONSE
ALLOGRAFT_REJECTION
REACTIVE_OXYGEN_SPECIES_PATHWAY
PANCREAS_BETA_CELLS
BILE_ACID_METABOLISM
PEROXISOME
NOTCH_SIGNALING
OXIDATIVE_PHOSPHORYLATION
MYC_TARGETS_V1
ADIPOGENESIS
DNA_REPAIR
ANDROGEN_RESPONSE
PROTEIN_SECRETION
APICAL_JUNCTION
SPERMATOGENESIS
XENOBIOTIC_METABOLISM
MTORC1_SIGNALING
FATTY_ACID_METABOLISM
GLYCOLYSIS
KRAS_SIGNALING_DN
TNFA_SIGNALING_VIA_NFKB
UV_RESPONSE_DN
TGF_BETA_SIGNALING
UV_RESPONSE_UP
HYPOXIA
ESTROGEN_RESPONSE_LATE
P53_PATHWAY
IL6_JAK_STAT3_SIGNALING
INFLAMMATORY_RESPONSE
APICAL_SURFACE
ESTROGEN_RESPONSE_EARLY
APOPTOSIS
EPITHELIAL_MESENCHYMAL_TRANSITION
IL2_STAT5_SIGNALING
KRAS_SIGNALING_UP
1 2 3 4 5 6 7 8 9 10 11 12 13 14 15
C
EPITHELIAL_MESENCHYMAL_TRANSITION
TNFA_SIGNALING_VIA_NFKB
ANGIOGENESIS
COAGULATION
APICAL_SURFACE
CHOLESTEROL_HOMEOSTASIS
NOTCH_SIGNALING
REACTIVE_OXYGEN_SPECIES_PATHWAY
COMPLEMENT
ESTROGEN_RESPONSE_EARLY
HEME_METABOLISM
SPERMATOGENESIS
MYOGENESIS
IL6_JAK_STAT3_SIGNALING
KRAS_SIGNALING_DN
APOPTOSIS
KRAS_SIGNALING_UP
HYPOXIA
TGF_BETA_SIGNALING
UV_RESPONSE_UP
INFLAMMATORY_RESPONSE
WNT_BETA_CATENIN_SIGNALING
ALLOGRAFT_REJECTION
G2M_CHECKPOINT
PROTEIN_SECRETION
ANDROGEN_RESPONSE
UV_RESPONSE_DN
IL2_STAT5_SIGNALING
MYC_TARGETS_V2
ESTROGEN_RESPONSE_LATE
E2F_TARGETS
HEDGEHOG_SIGNALING
APICAL_JUNCTION
FATTY_ACID_METABOLISM
MYC_TARGETS_V1
P53_PATHWAY
MITOTIC_SPINDLE
DNA_REPAIR
MTORC1_SIGNALING
OXIDATIVE_PHOSPHORYLATION
GLYCOLYSIS
PEROXISOME
PANCREAS_BETA_CELLS
UNFOLDED_PROTEIN_RESPONSE
XENOBIOTIC_METABOLISM
BILE_ACID_METABOLISM
INTERFERON_GAMMA_RESPONSE
PI3K_AKT_MTOR_SIGNALING
ADIPOGENESIS
INTERFERON_ALPHA_RESPONSE
13 14 15

**Figure S5. Patterns of Hallmark pathway enrichment analysis.** A) Heatmap showed the two-way hierarchical clustering of enrichment scores in each pathway. X-axis is the cell cluster id. The redder the color, the higher of enrichment scores. Numbers in the left-side of the heatmap showed the defined patterns. B) Heatmap showed the row hierarchical clustering of enrichment scores in each pathway. C) Heatmap showed the two-way hierarchical clustering of enrichment scores in IgM dominant cell lineage using DEGs are specifically differentially expressed in IgM lineage compared with DEGs detected by comparisons made between IgG1 dominant clusters.

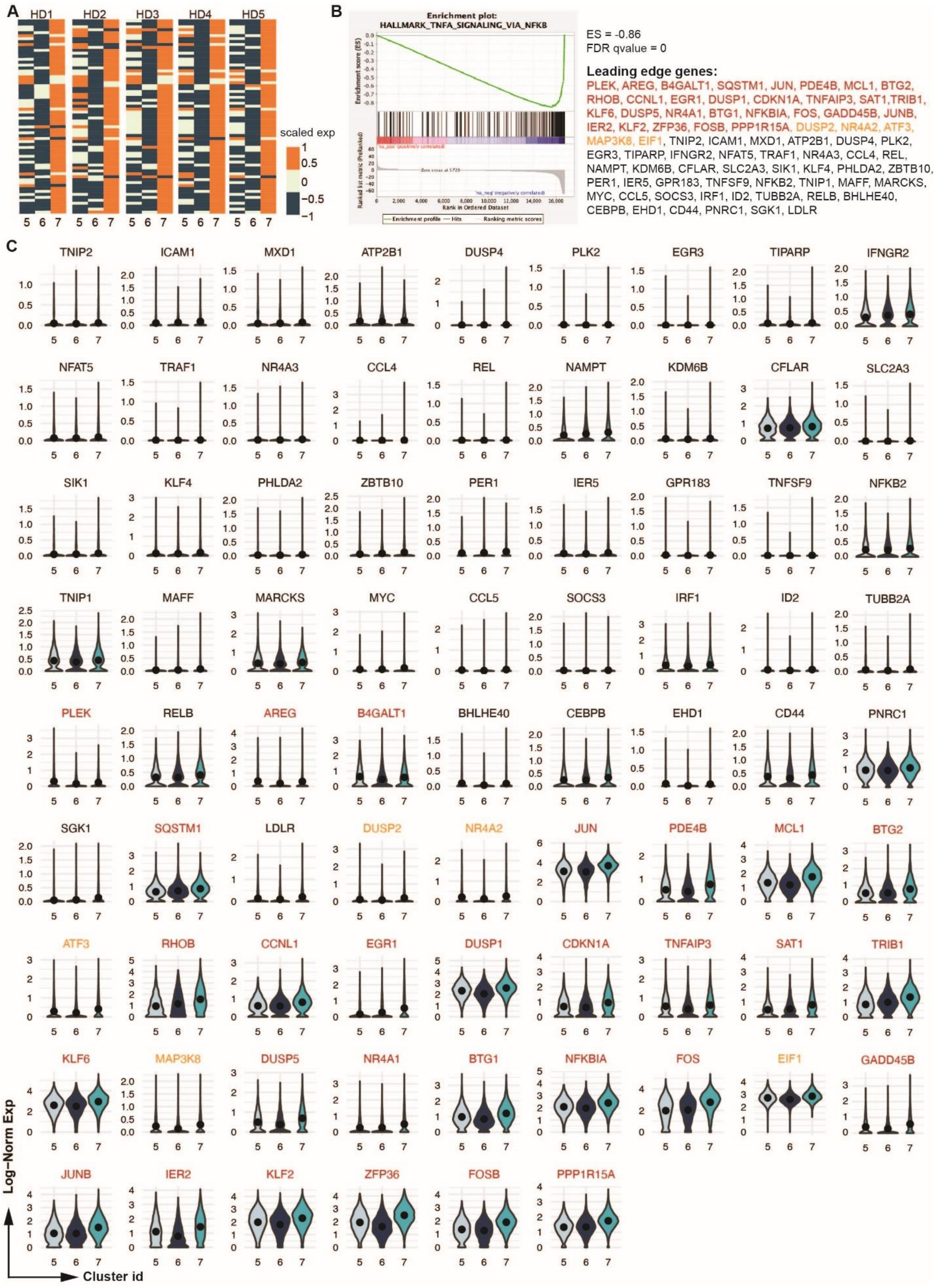
A
HD1
HD2
HD3
HD4
HD5
scaled exp
1
0.5
0
−0.5
−1
5 6 7
B
Enrichment plot:
HALLMARK_TNFA_SIGNALING_VIA_NFKB
Enrichment score (ES)
Ranked list metric (PreRanked)
Rank in Ordered Dataset
Enrichment profile
Hits
Ranking metric scores
ES = -0.86
FDR qvalue = 0
Leading edge genes:
PLEK, AREG, B4GALT1, SQSTM1, JUN, PDE4B, MCL1, BTG2, RHOB, CCNL1, EGR1, DUSP1, CDKN1A, TNFAIP3, SAT1,TRIB1, KLF6, DUSP5, NR4A1, BTG1, NFKBIA, FOS, GADD45B, JUNB, IER2, KLF2, ZFP36, FOSB, PPP1R15A, DUSP2, NR4A2, ATF3, MAP3K8, EIF1, TNIP2, ICAM1, MXD1, ATP2B1, DUSP4, PLK2, EGR3, TIPARP, IFNGR2, NFAT5, TRAF1, NR4A3, CCL4, REL, NAMPT, KDM6B, CFLAR, SLC2A3, SIK1, KLF4, PHLDA2, ZBTB10, PER1, IER5, GPR183, TNFSF9, NFKB2, TNIP1, MAFF, MARCKS, MYC, CCL5, SOCS3, IRF1, ID2, TUBB2A, RELB, BHLHE40, CEBPB, EHD1, CD44, PNRC1, SGK1, LDLR
C
TNIP2
ICAM1
MXD1
ATP2B1
DUSP4
PLK2
EGR3
TIPARP
IFNGR2
NFAT5
TRAF1
NR4A3
CCL4
REL
NAMPT
KDM6B
CFLAR
SLC2A3
SIK1
KLF4
PHLDA2
ZBTB10
PER1
IER5
GPR183
TNFSF9
NFKB2
TNIP1
MAFF
MARCKS
MYC
CCL5
SOCS3
IRF1
ID2
TUBB2A
PLEK
RELB
AREG
B4GALT1
BHLHE40
CEBPB
EHD1
CD44
PNRC1
SGK1
SQSTM1
LDLR
DUSP2
NR4A2
JUN
PDE4B
MCL1
BTG2
ATF3
RHOB
CCNL1
EGR1
DUSP1
CDKN1A
TNFAIP3
SAT1
TRIB1
KLF6
MAP3K8
DUSP5
NR4A1
BTG1
NFKBIA
FOS
EIF1
GADD45B
JUNB
IER2
KLF2
ZFP36
FOSB
PPP1R15A
Log−Norm Exp
Cluster id

**Figure S6. Genes driven the difference of TNFα signaling via NFKB pathway in late LLPC.**

A) Heatmap showed the scaled corresponding TNFα singaling via NFκB pathway enriched maturation-associated DEG expressions in cluster 5, 6 and 7 in each individual. B) GSEA prerank analysis results using preranked genes between cluster 6vs7 (left); enrichment score, FDR qvalue, and leading-edge genes were listed (right). Gene colored in red were DEGs from Table S4 and in orange were cluster 6vs7 specific DEGs included in Table S6. C) Expression of differentially expressed leading-edge genes (|avgLogFC| > 0.25, Bonferroni adjusted p value < 0.05) from (B) in cluster 5, 6 and 7, the order of genes was corresponding to the gene rank in preranked gene list in Table S6.

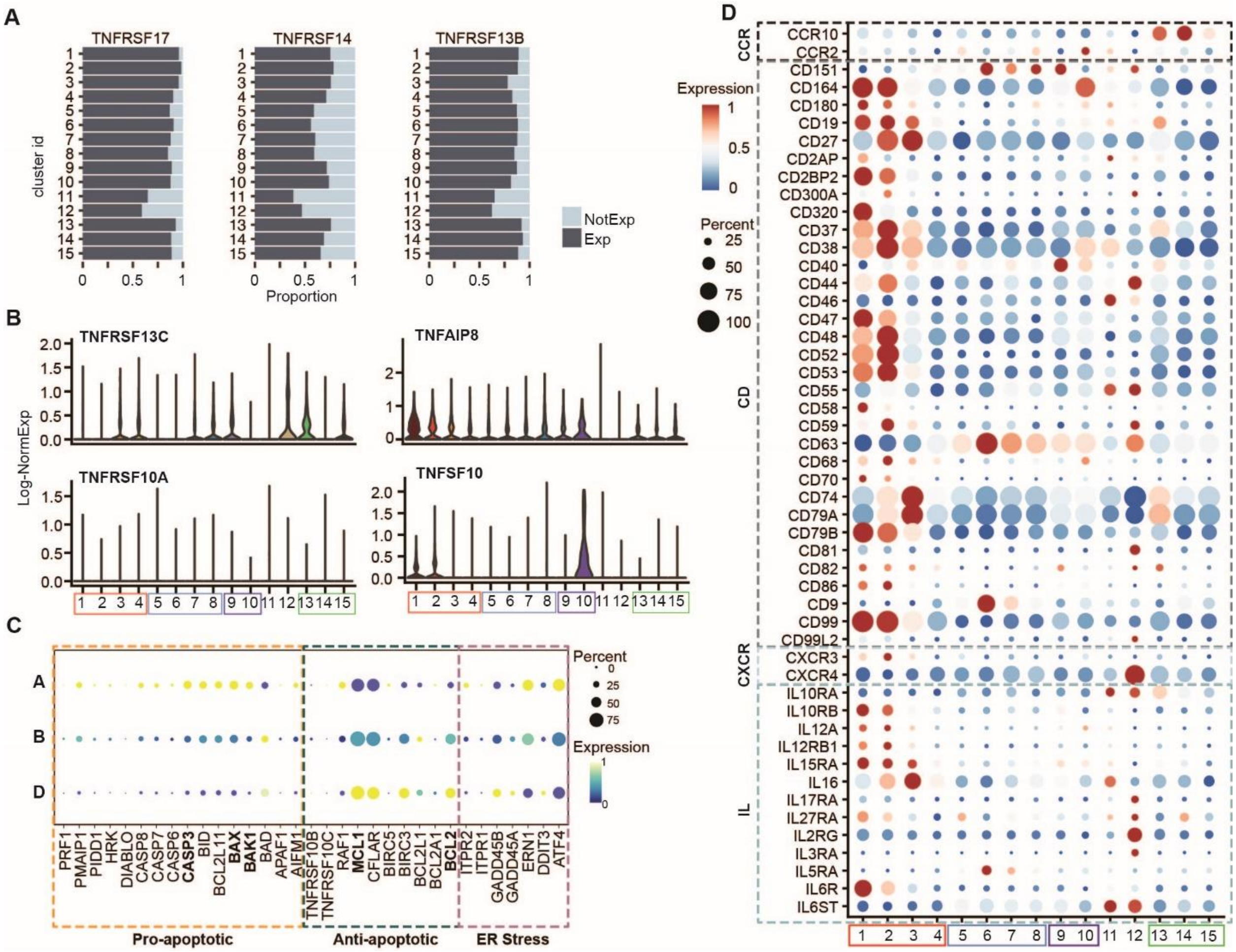


**Figure S7. Visualization of indicative genes**. A) Light and dark blue bar plots showed the proportion of cells in each cluster not expressed (NotExp) and expressed (Exp) gene TNFRSF17, TNFRSF14 and TNFRSF13B. B) Violin plots showed the expression of remaining expressed TNF family genes. C) Dotplot visualized the gene expression of pro-apoptotic, anti-apoptotic and ER-stress associated genes based on the FACS-sorted cell labels. The bluer the dots, the lower the gene expression. The size of dots represented the percentage of cells from associated cell population expressed the corresponding genes. D) Dotplot showed the gene expression from CCR, CD, CXCR ad interleukin (IL) families.

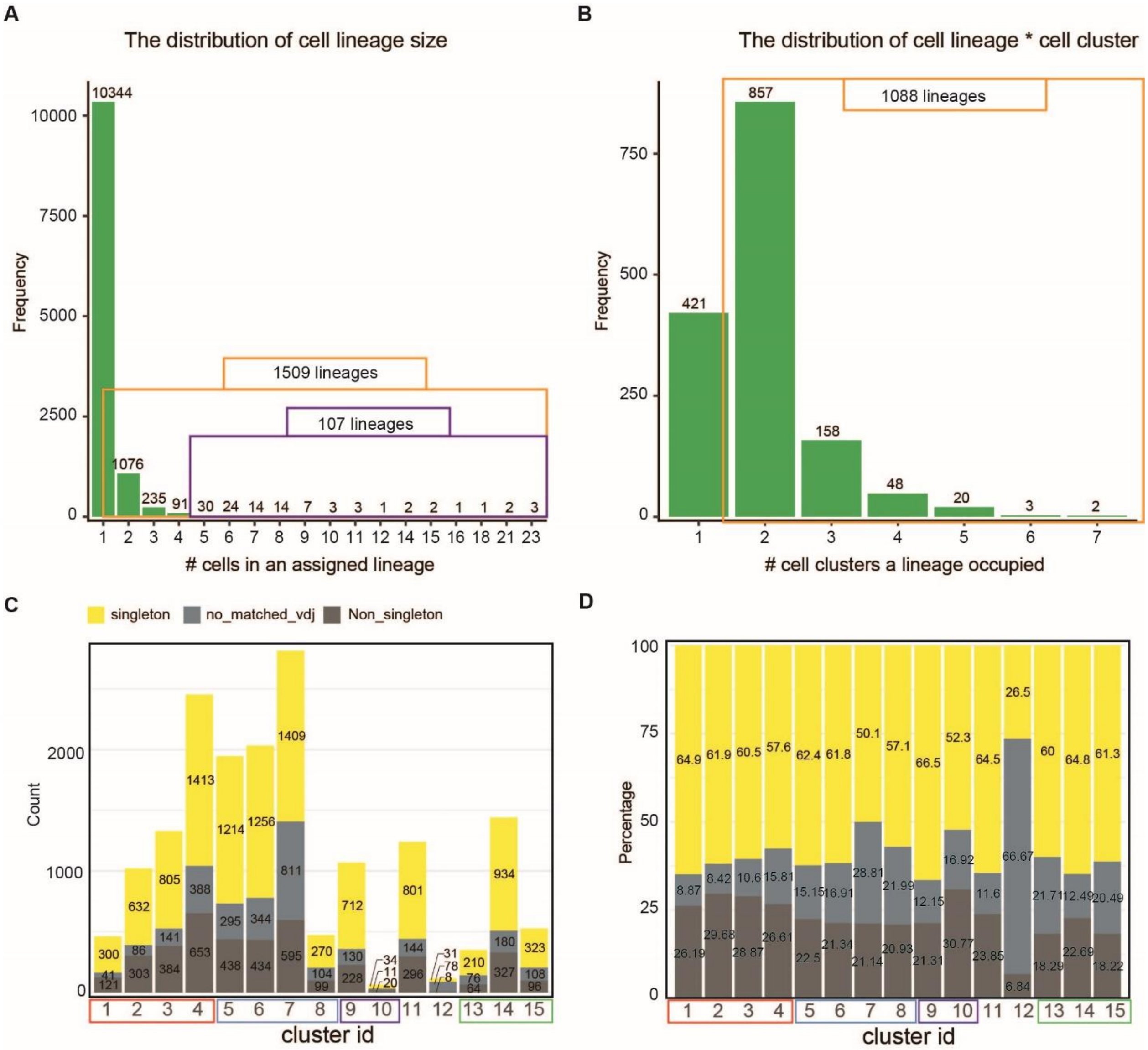


**Figure S8. Summary of scVDJ-seq data detected clones/lineages.** A) The distribution of lineage size. X-axis is the number of cells contained in detected cell lineages. Y-axis is the number of cell lineages. Orange and purple boxes summarize the total number of lineages that have at least 2 or 5 cells. B) The distribution of overlapped cell clusters of each non-singleton lineages. Orange boxes summarize the number of lineages having at least 2 cells and overlapped with at least 2 cell clusters. C) and D) Count the total number (C) and percentage (D) of cells in each cell cluster by if cells have matched scVDJ-seq information, if they are from singleton lineages (yellow) or not (dark grey). Cells with no matched scVDJ information were colored in ash.

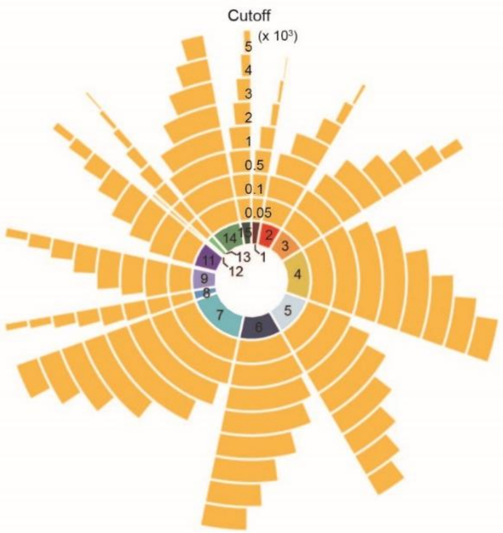


| cluster_id | count | 50 | 100 | 500 | 1000 | 2000 | 3000 | 4000 | 5000 |
|---|---|---|---|---|---|---|---|---|---|
| 1 | 462 | 99.6 | 97.6 | 86.8 | 66.9 | 31.2 | 16.7 | 8.9 | 3 |
| 2 | 1021 | 99.8 | 99.3 | 87.3 | 66.4 | 33.3 | 15.8 | 6.7 | 2.5 |
| 3 | 1330 | 99.2 | 98 | 87.1 | 74.3 | 49.9 | 33.6 | 20.1 | 11.7 |
| 4 | 2454 | 99.9 | 99.5 | 92 | 79.6 | 58.5 | 41.4 | 29.5 | 20.6 |
| 5 | 1947 | 99.4 | 98.8 | 88.5 | 76.8 | 55 | 39 | 27.7 | 19.3 |
| 6 | 2034 | 99.5 | 99.1 | 94.4 | 83.3 | 63.9 | 46.8 | 35.1 | 26.2 |
| 7 | 2815 | 99.6 | 99.3 | 91.5 | 79.1 | 53.9 | 36.1 | 23.8 | 15.1 |
| 8 | 473 | 98.9 | 98.1 | 90.1 | 79.5 | 58.4 | 42.1 | 27.5 | 20.9 |
| 9 | 1070 | 99.6 | 99.2 | 87.4 | 72.2 | 45.6 | 27.9 | 18.2 | 11.3 |
| 10 | 65 | 96.9 | 96.9 | 89.2 | 76.9 | 52.3 | 38.5 | 18.5 | 13.8 |
| 11 | 1241 | 98 | 94.8 | 76.1 | 57.9 | 36.8 | 23.5 | 14.6 | 9.2 |
| 12 | 117 | 100 | 99.1 | 72.6 | 40.2 | 6.8 | 0.9 | 0 | 0 |
| 13 | 350 | 99.7 | 99.7 | 97.7 | 83.7 | 59.1 | 36.9 | 24 | 12.6 |
| 14 | 1441 | 99.7 | 99.4 | 95.2 | 85.7 | 64.1 | 45.2 | 29.8 | 18.9 |
| 15 | 527 | 100 | 100 | 93 | 80.3 | 56.9 | 41.7 | 26.6 | 18 |

**Figure S9. Summary of Ig heavy chain gene transcript numbers in each cell cluster.** Circos plot summarizes the proportion of cells in each cell cluster that pass the criterion by using different cutoffs to gate Ig heavy chain genes. The most inner circle is colored as Fig. 1C and numbers indicate corresponding cell cluster id. From inner to outer circle, the cutoffs are 50, 100, 500, 1000, 2000, 3000, 4000 and 5000. The table summarizes the percentage of cells in each cell cluster that passed cutoffs. The count column shows the total number of cells in each cell cluster.

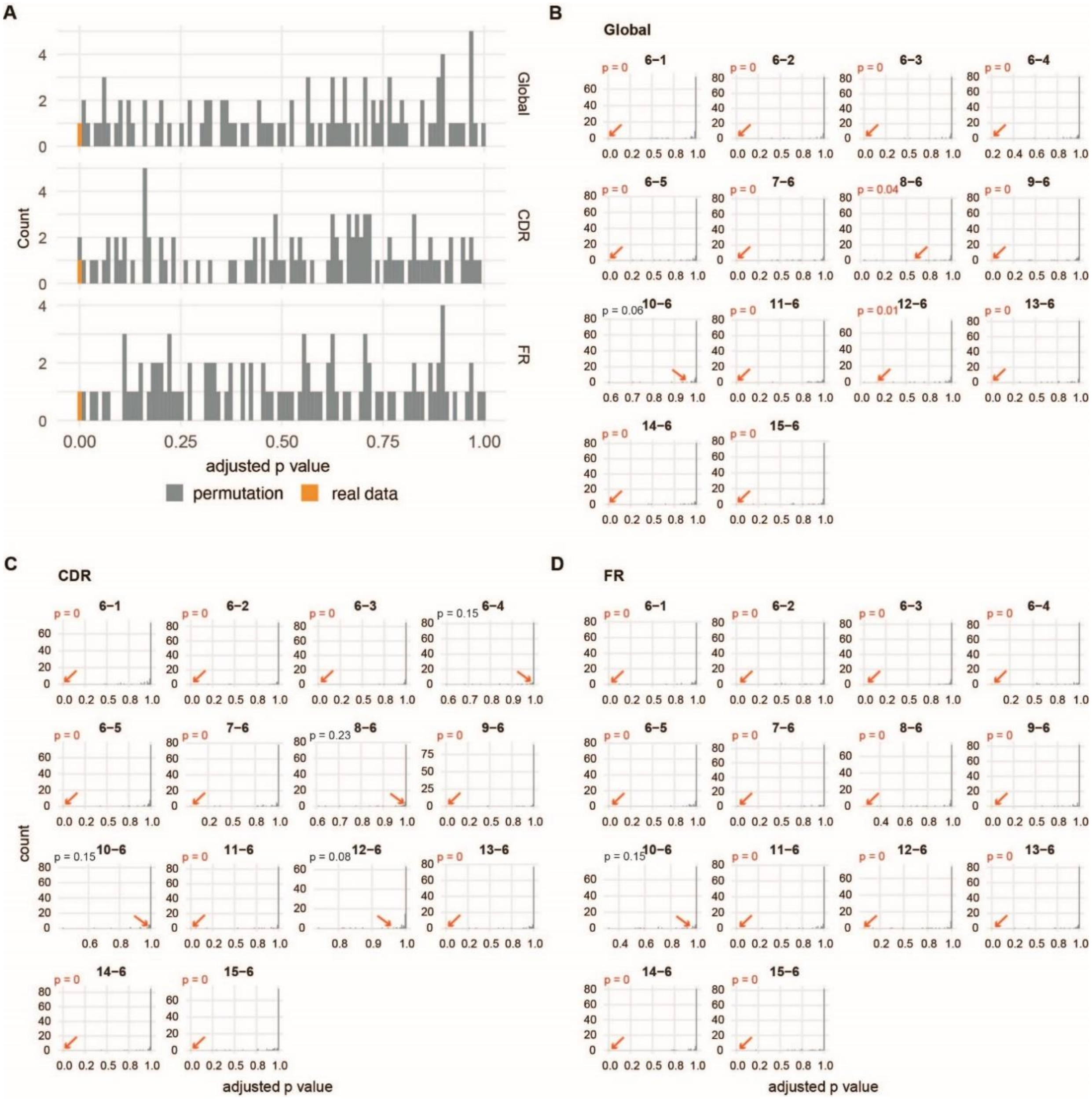


**Figure S10. Permutation results for the mutation frequency test.** A) Barplot showing the distribution of p values from ANOVA test for the global heavy chain variable region (Global), CDR and FR mutation frequency using real data and shuffled data from permutation test. Orange bars and grey bars indicated the p values from using real and randomly shuffled datasets. B) – D) The distribution of p values from Tukey's HSD (honestly significant difference) tests for the Global (B), CDR (C) and FR (D) mutation frequency between cluster 6 and other 14 clusters by using real and randomly shuffled datasets. Red arrows pointed out the adjusted p value from Tukey's HSD test using real data, left upper side showed the p value from permutation test and significant results were colored in red. Titles indicated the objects of comparison.

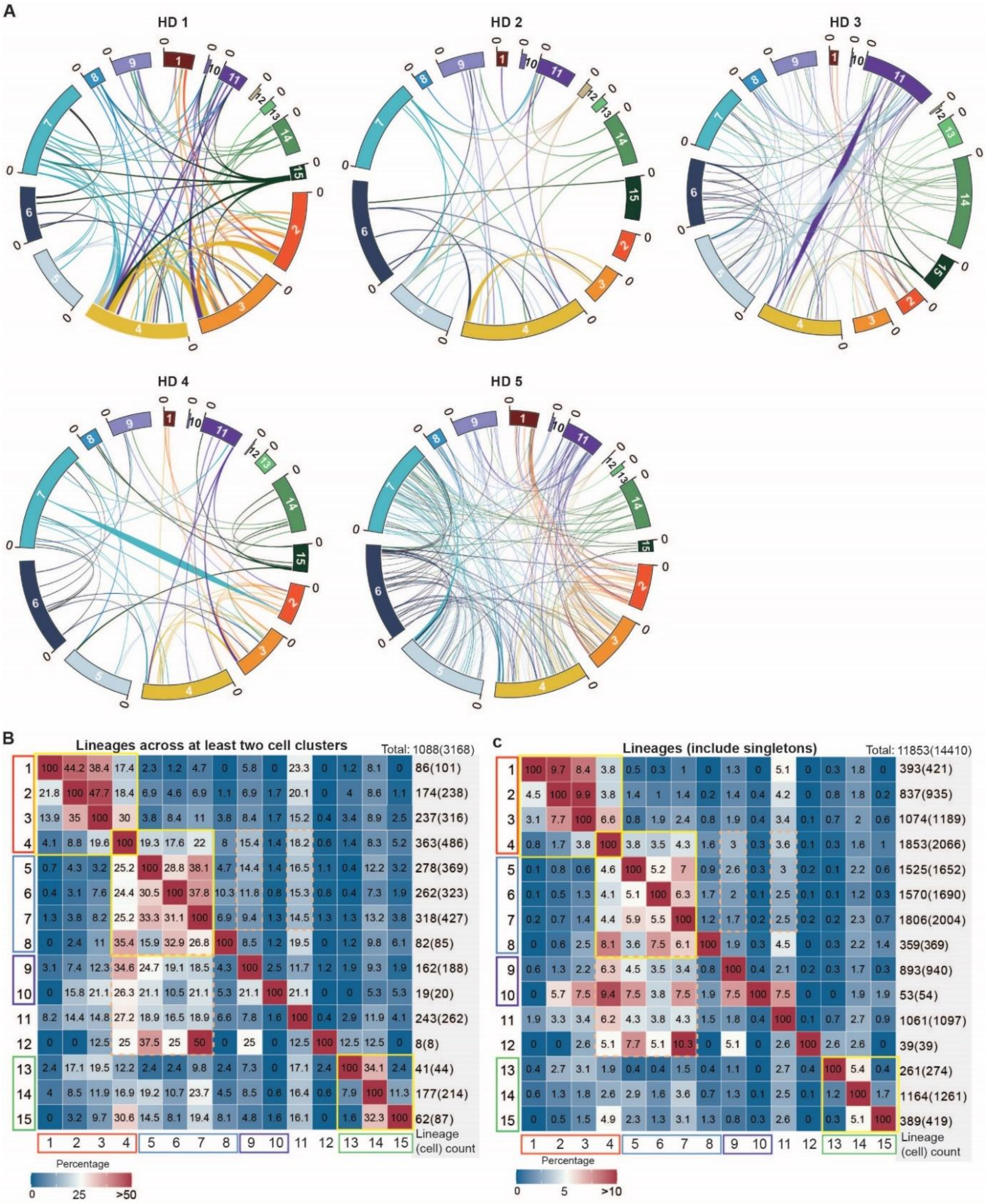

**Figure S11. Clonal connectivity visualization in all cell clusters.** A) Circos plot from main IgG1 dominant cell clusters 1 to 8 in a connecting detected lineages in each donor. Lineages were colored by the latest cell cluster; lineages spanned in both path 1 (c6) and path 2 (c7 and/or c8) were colored in purple, indicating late phase. B) – C) Percentage of connected clones between any two cell clusters when only considering lineages across at least two cell clusters (B) or lineages include singletons (C). Numbers in each block showed the percentage of lineages from each row cluster that were connected with other. The numbers showing in the grey boxes recorded the total number of lineages and cells showing in bracket in associated cell cluster. Yellow boxes highlighted the highly connected cell blocks and dashed orange box highlighted those clones showing high connectivity in one direction but not in another.

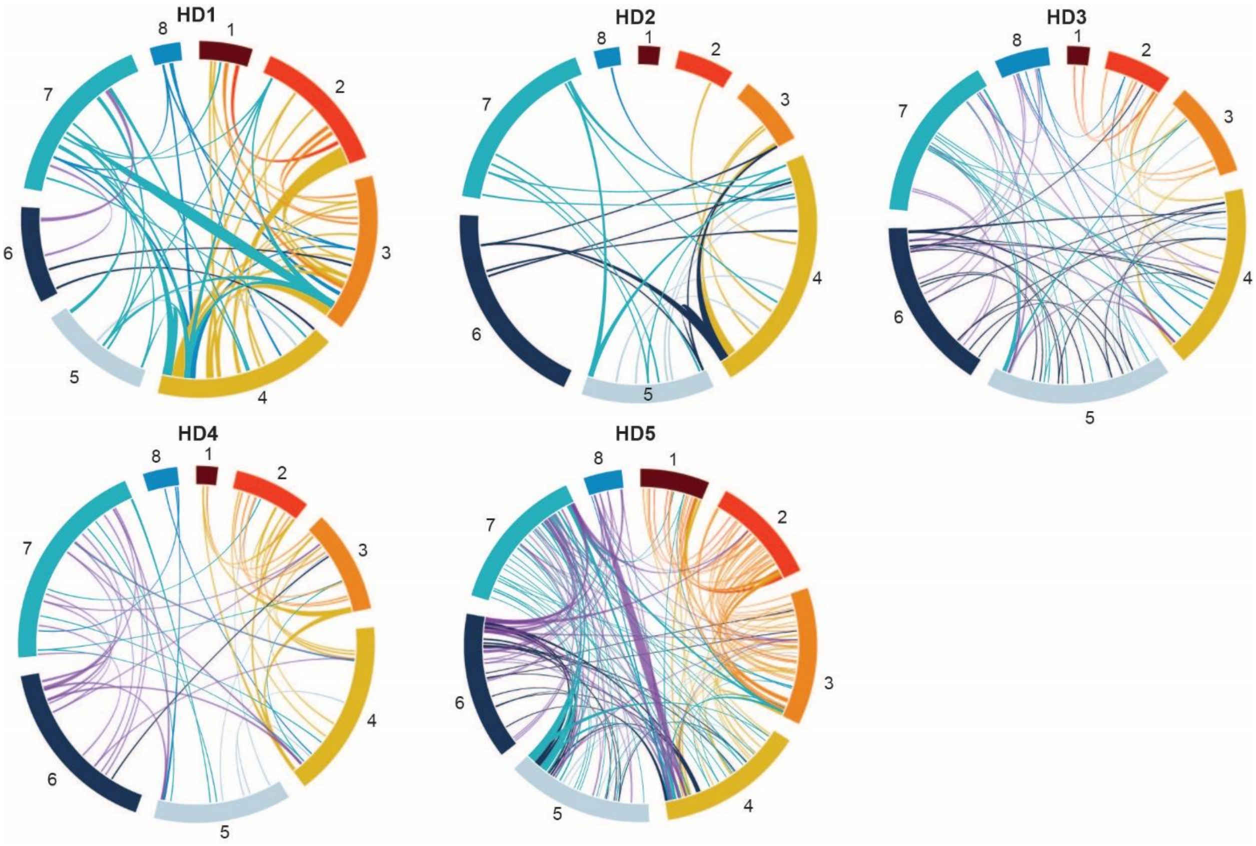

**Figure S12. Clonal connectivity of nearly identical clones in IgG1-dominant clusters 1 to 8.** A homology of 98% was used within the CDR3 region to cluster clones within each of the 5 subjects. Cell clusters are represented by the outer ring colors and sequences are grouped into clones in size descending order from counterclockwise to clockwise. Lines connecting cell clusters indicate a clone that was found in multiple clusters.
